# DCN1 inhibitor induces fetal hemoglobin through self-limited regulation of CUL3 neddylation

**DOI:** 10.64898/2026.04.22.720177

**Authors:** Samuel A. Miller, Anthony Monti, Enhua H. Zhou, Chi Hua Sarah Lin, Orna Lynch, Sayani Mukherjee, Cassie Hei, Robert Nicewonger, Colin Diner, Alejandra Macias-Garcia, Kristen Marino, Na Li, Ravi Ujjinamatada, Akshay Salegaonkar, Sunil Kuppasani, Max Rogers-Grazado, Corey Hayford, Frank Lovering, Qing Tang, Yihua E. Ye, Wei Zhang, Timothy Fulton, Jennifer J. O’Brien, Xiaoji Sun, Atli Thorarensen, Mauricio Cortes, Cameron C. Trenor, Sriram Krishnamoorthy

## Abstract

Few genetic loci are as well-characterized as the globin gene locus, and the substitution of healthy γ-globin (HbF) for missing or mutated β-globin (HbB) is an established therapeutic strategy for β-hemoglobinopathies including sickle cell disease (SCD) and β-thalassemia. Although substantial progress has been made in understanding HbF derepression and globin switching, many current therapeutic strategies involving small molecules increase HbF through broad epigenetic perturbations or cytotoxic stress, raising concerns about dose-limiting cytopenias and off-target effects. By coupling single-cell transcriptomics, genetic perturbations, and functional genomics, we identified an unknown role of neddylation in the regulation of fetal hemoglobin (HbF). Partial impairment of neddylation of cullin ubiquitin ligase 3 (CUL3) through defective in cullin neddylation 1 (DCN1) inhibition leads to highly selective chromatin changes, histone demethylation, and globin locus binding of known activators of HbF transcription. Further, DCN1 inhibition drives globin switching and HbF increases *in vitro* and *in vivo* with minimal off-target transcriptional effects and no evidence of cytotoxicity or stress erythropoiesis. To therapeutically target this axis, we report the discovery and characterization of CLY-124, a first-in-class, covalent DCN1 inhibitor with favorable pharmacology properties. In a humanized mouse model, CLY-124 showed a dose-dependent increase in HbF as monotherapy and in synergy with hydroxyurea (HU), a current standard of care. Collectively, these findings highlight the power of single-cell transcriptomics to elucidate undiscovered biologic insights with therapeutic potential, and the promise of DCN-1 inhibitors like CLY-124 to address β-hemoglobinopathies. With an appropriate nonclinical safety profile, a first-in human study of safety, pharmacokinetics and HbF assessments in healthy volunteers and participants with SCD is ongoing for CLY-124.

**One-sentence Summary:** DCN1 is a promising target for ß-hemoglobinopathies

## Introduction

Sickle cell disease (SCD), a β-hemoglobinopathy, is a common inherited monogenic disorder caused by a substitution mutation in the β-globin gene (HBB), which leads to hemoglobin polymerization upon deoxygenation, affecting the health and function of red blood cells (*1*). The resulting downstream sequelae include rapid hemolysis, chronic anemia, and recurrent vaso-occlusive pain crisis (*2*). Increasing fetal hemoglobin (HbF) is a well-established therapeutic goal for SCD which ameliorates these symptoms (*3*). Human genetics has validated HbF as safe and disease correcting in SCD, as exemplified by hereditary persistence of fetal hemoglobin (HPFH), where there is incomplete fetal hemoglobin gene (HBG1/2) repression in adults (*4*). Moreover, high HbF levels have been linked with lower childhood mortality and increase in life expectancy in patients with SCD (*5*).

Intricate molecular pathways orchestrate the temporal regulation of HBB and HBG1/2 during fetal and adult stages of human life. The interplay of multiple genetic regulators is instrumental in the expression of HBG1/2 in the fetus and repression in adult life leading to its silencing and concurrent expression of HBB (*6*). Available therapies for SCD, as well as other β-hemoglobinopathies, including those designed to de-repress HBG1/2; all have various limitations, leaving substantial unmet need for better differentiated safety profiles, more favorable risk-to-benefit ratios, increased convenience for patients, and increased accessibility (*7*). The current standard of care, hydroxyurea (HU) approved after pivotal Multicenter Study of Hydroxyurea (MSH) trials (*8*), exerts its beneficial effects primarily through the induction of HbF (*9*). However, its clinical efficacy is limited by dose-dependent side effects such as myelosuppression, neutropenia, and reduced reticulocyte counts (*10*) and the response to treatment is widely variable (*11*). It is recommended that HU may be combined with agents that activate HBG transcription to achieve greater HbF levels thereby improving efficacy (*12*). No small molecule has demonstrated clinical superiority over HU without treatment emergent adverse effects, and no small molecule combination with HU has demonstrated an adverse event-free safety profile in clinically tested dosing regimens (*13*). Therefore, identifying clinical agents that can safely boost the efficacy of HU or replace it are urgently needed (*12*). Small molecules have been identified that induce HbF via stress erythropoiesis-associated mechanisms, but are accompanied by cytotoxicity and cytopenias, limiting their therapeutic utility (*14*). Small-molecule clinical candidates have also been developed to directly disrupt chromatin-regulating complexes. Inhibitors directed to these complexes, though efficacious, have shown adverse effects in tested SCD patients (*15*). Newer approaches targeting different epigenetic regulators such as Wiz, EED (a component of PRC2), and ZBTB7A are currently in clinical development. Since these complexes regulate numerous other genomic loci, their inhibition can broadly affect diverse biological processes beyond the expression of fetal hemoglobin (*16*).

Recently, an autologous cell-based gene therapy targeting the enhancer of BCL11A (*17*), a key repressor of HBG1/2 (*18*), was approved to increase HbF levels (*18*), focusing on selective modulation of fetal hemoglobin alone without extensively perturbing genomic loci. Despite their promises, access to these therapies remains limited due to its complexity, high cost, and high risk of toxicity to patients. Collectively, these shortcomings have spurred efforts to discover safer and more tractable approaches to HbF induction (*7*).

Phenotypic screens and target-based drug discovery campaigns for HbF induction have historically relied on cell lines such as K562 erythroleukemia cells and human umbilical cord blood-derived erythroid progenitor 2 cells (HUDEP-2), which are already committed to adult globin expression (*19*) (*20*) (*20*). While these systems are experimentally tractable and scalable, they are biased toward identifying mechanisms that derepress silenced HBG1/2 in late erythroid cells and may miss regulators that act earlier in hematopoietic progenitors (*21*). These early progenitors could reveal targetable biology driving physiological HBG1/2 expression, rather than relying on de-repression of already-silenced genes. Primary human hematopoietic cells offer a potential solution by enabling discovery in a more developmentally relevant context. However, the added complexity of culturing and differentiating these cells limits assay scalability thereby restricting the chemical space that can be explored. These considerations reflect a broader need for generalizable discovery frameworks capable of efficiently and effectively linking relevant cell states to actionable therapeutic hypotheses across areas of unmet clinical need.

In this context, we recently described a lab-in-the-loop platform that leverages high-throughput transcriptomics as a proxy for cell state within an active-learning framework, enabling the prediction and prioritization of compounds for phenotypic testing (*22*). Here, we applied this approach to sickle cell disease (SCD), using single-cell transcriptomic profiling of erythropoiesis in stem cells and early progenitors, prior to lineage commitment, to uncover previously unrecognized mechanisms of HbF regulation. Integrated analyses of screening hits implicated neddylation in globin gene regulation. Mechanistic interrogation of this pathway revealed a central constraint: broad disruption of CUL3 induces HbF but impairs erythroid differentiation, whereas partial suppression of CUL3, up to ∼60%, preserves erythropoiesis while retaining HbF induction. We found that Defective in Cullin Neddylation 1 (DCN1), a positive regulator of CUL3 neddylation, resolves this constraint because its perturbation is intrinsically self-limited, where even maximal pharmacologic or genetic inhibition of DCN1 did not suppress CUL3 neddylation beyond the threshold compatible with erythropoiesis. Through mechanistic deconvolution and structure-guided chemical optimization, we established DCN1 as a target for HbF induction with no detectable adverse effects on hematopoiesis or cell viability and developed CLY-124, a first-in-class DCN1 inhibitor. CLY-124 outperformed HU in inducing HbF in a humanized mouse model and showed synergistic activity with HU even at low doses, supporting DCN1 inhibition as a therapeutic strategy for SCD and other β-hemoglobinopathies. CLY-124 has now entered early phase clinical development in healthy volunteers and patients with SCD.

## RESULTS

### Complete inhibition of CUL3 upregulates fetal hemoglobin but impairs erythropoiesis

To discover HbF inducers, we conducted a drug screen informed by single-cell transcriptomic data spanning human erythropoiesis. We focused on identifying mechanisms acting before HSPC commitment to adult or fetal erythroid fates, rather than altering HBG expression in committed cells. Therefore, primary human CD34^+^ cells were perturbed at the HSPC stage, and HBG induction was assessed in erythroblasts (**fig. S1A**). This screen identified compounds with diverse mechanisms, including known but clinically limited pathways such as histone deacetylase and cell cycle inhibition, as well as previously uncharacterized mechanisms, notably neddylation inhibition. Treating primary CD34^+^ cell-derived erythroid cells with screening hit compound, MLN4924, a well-characterized inhibitor of NEDD8-activating enzyme E1 (NAE1) (*23*), led to a dose-dependent increase in fetal hemoglobin, quantified at differentiation day 14 by HPLC as the percentage of total hemoglobin (%HbF) (**Fig. 1A**), however cytotoxicity was observed in the highest tested doses, 0.1 μM and above (**fig. S1B**).

**Fig. 1.**
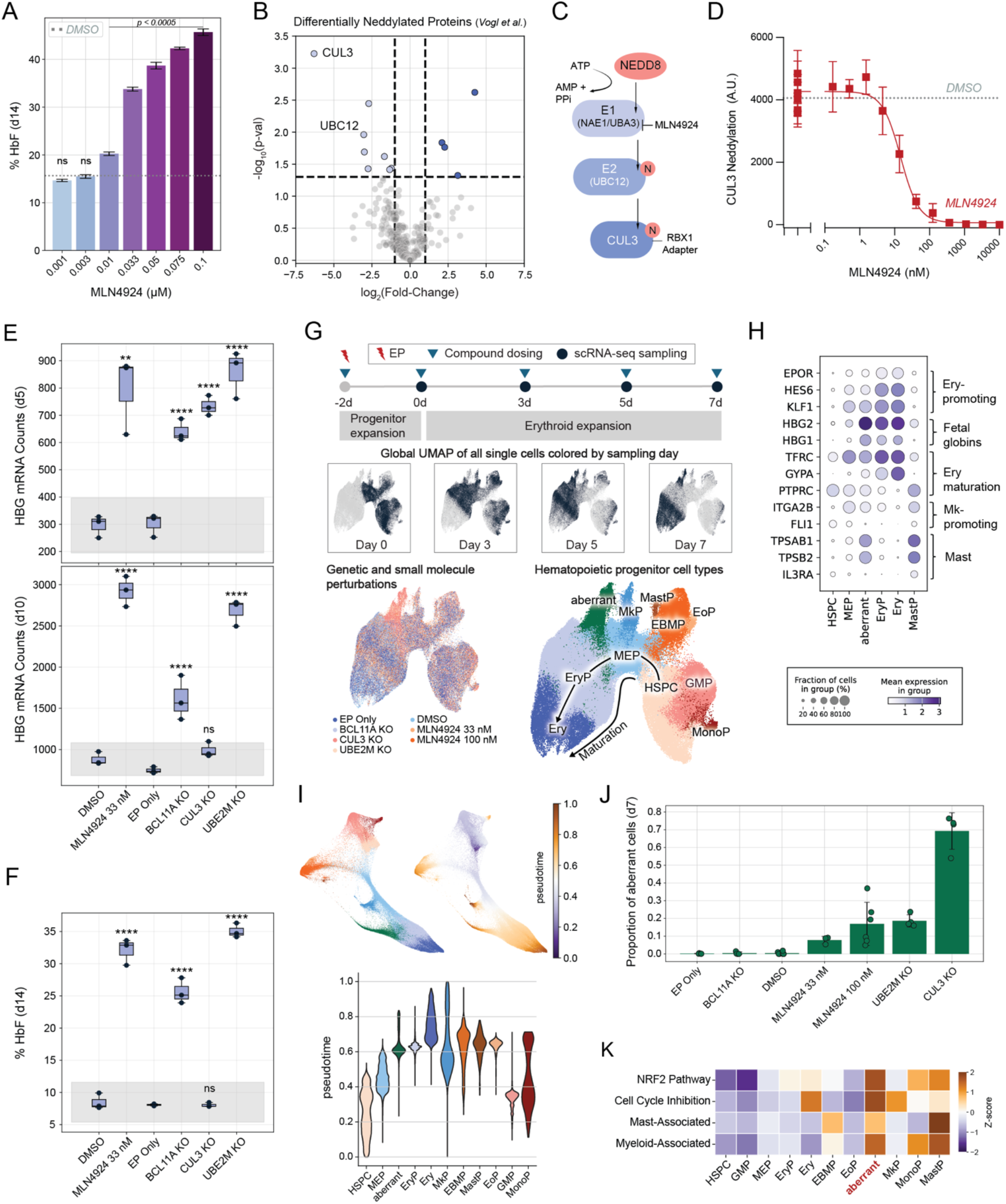
Modulating CUL3 activity induces fetal hemoglobin but disrupts erythropoiesis. **(A)** Dose-response curve of MLN4924 showing % HbF relative to total hemoglobin on day 14, measured by HPLC. Statistical significance was assessed versus DMSO using one-way ANOVA with Dunnett’s multiple comparisons test (N=3). **(B)** Differentially neddylated proteins identified by sNUSP mass spectrometry following MLN4924 treatment in HEK293 cells. **(C)** Schematic illustrating NEDD8 transfer from E1 to E2 to the CUL3-containing E3 ligase complex. **(D)** Inhibitory dose-response of MLN4924 against CUL3 neddylation in HUDEP2 cells, measured by AlphaLISA (N=4). **(E)** Normalized HBG expression on days 5 and 10 measured by nCounter. (**F**) % HbF at day 14 quantified by HPLC. In (E) and (F), statistical significance was determined using unpaired t-test or one-way ANOVA with Dunnett’s multiple comparisons test, as appropriate (**P<0.005, ****p<0.00005). Grey box indicates 95% confidence interval computed across control conditions (DMSO and EP Only). **(G)** Schematic and representative UMAP embeddings from time course scRNA-seq following compound and CRISPR perturbations. **(H)** Gene expression profiles across selected cell types from scRNA-seq. **(I)** FA2 embeddings showing cell types and inferred pseudotime, with accompanying violin plots of pseudotime distributions across cell types. **(J)** Proportion of aberrant cells on day 7 from scRNA-seq. **(K)** Heatmap of Z-scored pathway signature enrichment across cell types. Panels (A) and (E-K) were performed in mPB CD34^+^ cells.

To identify neddylation pathway components downstream of NAE1 that may contribute to HbF induction, we analyzed a publicly available dataset of proteome-wide changes in neddylation following NAE1 inhibition in HEK293 cells (*24*). This analysis identified CUL3 and UBC12 (*UBE2M*) among the proteins showing the greatest reductions in neddylation after NAE1 inhibition (**Fig. 1B**). NAE1 is a subunit of the heterodimeric UBA3-NAE1 E1 enzyme that activates NEDD8 and transfers it to the catalytic cysteine of the E2 enzyme UBC12 (*25*) (**Fig. 1C**). UBC12 subsequently mediates NEDD8 conjugation to cullin family proteins, including CUL3, a modification required for activation of CUL3-containing cullin-RING ligase complexes, which promote ubiquitination of substrate proteins and thereby regulate their abundance and function (*26*) (*24*). We therefore used CUL3 neddylation as a proximal functional readout of pathway inhibition and confirmed that MLN4924 suppressed CUL3 neddylation in a dose-dependent manner in HUDEP-2 cells, with an IC_50_ of 14.87 nM and maximal inhibition (E_max_) of ∼100% (**Fig. 1D**).

Structural studies have demonstrated CUL3 functions as a scaffold protein in a subclass of E3 ubiquitin ligases, bridging BTB-domain-containing substrate adaptors with the RING finger protein RBX1 (*27, 28*) (**Fig. 1C**). These studies further suggest that BTB-domain-containing adaptors form homodimers, each engaging one CUL3 molecule, and that CUL3 neddylation induces a conformational change that promotes efficient ubiquitin transfer. To assess the effects of disrupting this pathway during erythroid differentiation, we quantified HBG expression and HbF levels across a 14-day differentiation time course using the BCL11A enhancer knockout as a positive control. CRISPR knockout of either UBC12 or CUL3 initially phenocopied the HBG induction observed with MLN4924 treatment, supporting a mechanistic role for this pathway in regulating HBG expression during erythroid differentiation (**Fig. 1E and fig. S1C-E**). However, the effect of CUL3 KO was transient: *HBG* expression was increased at day 5 but not at day 10 relative to EP only control (**Fig. 1E**), with no corresponding increase in HbF protein at day 14 (**Fig. 1F**). In contrast, UBE2M KO produced sustained *HBG* induction through day 10 and increased %HbF at day 14 to levels comparable to those achieved with MLN4924 (**Fig. 1E,F**).

To further investigate why CUL3 KO failed to induce comparable levels of HBG transcript expression on day 10, we profiled these genetic and small molecule perturbations in CD34^+^ cells over erythroid differentiation time using scRNA-seq on days 0, 3, 5, and 7 (**Fig. 1G and fig. S1E,F**). Beginning on differentiation day 3, perturbed conditions gave rise to an aberrant cell state that showed no evidence of technical abnormality by standard QC metrics (**fig. S1E and fig. S2A**) and was characterized by a megakaryocyte-erythroid progenitor (MEP)-like transcriptional profile with HBG expression together with select functional mast cell markers (**Fig. 1H and fig. S1G**).

Pseudotime ordering of sequenced cells across all timepoints (**Methods**) revealed that at the terminal stage of MEP maturation, cells split into two trajectories-either an aberrant lineage or canonical erythropoiesis (**Fig. 1I**). CUL3 KO led to the most dramatic enrichment of cells in the aberrant state where 69% of progenitors were aberrant on differentiation day 7 (**Fig. 1J and fig. S2A,B**). A dose-dependent effect was observed in MLN4924 treated cells where 0.1µM resulted in a similar proportion of cells in the aberrant trajectory as UBE2M KO at day 7, in all cases reducing the fraction of committed erythroid progenitors. Further characterization of the aberrant cell population revealed enrichment of NRF2 activation genes (*NQO1, TXNRD1, SQSTM1, FTH1, FTL, TXN, GSTO1*) consistent with the role of CUL3 as a negative regulator of this pathway (*29*) (**Fig. 1K**). We additionally observed enrichment of cell cycle inhibitor gene expression (*CDKN1A, CDKN1B, CDKN2A, CDKN2B*) which was accompanied by an elevated number of cells inferred to be in G0 and G1 phase of cell cycle compared to committed erythroid progenitors, inferred to be cycling in G2 and S phase (**Fig. 1K and fig. S2C-E**).

As noted above, CUL3 KO induced robust *HBG* expression at differentiation day 5, but this response diminished by day 10. scRNA-seq further showed that CUL3 KO MEPs were diverted into an aberrant cell state rather than continuing along the erythroid differentiation trajectory. Together, these findings suggest that HBG upregulation is initiated before erythroid commitment but is not sustained because the cells are redirected toward an alternate fate and thus may fail to complete productive erythropoiesis. Consistent with this model, we observed upregulation of non-erythroid transcriptional programs, including myeloid-associated genes (*CSF1, APOC1, CST3, S100A10, LAPTM5, CCRL2, TRPV2*) and mast-associated genes (*KIT, TPSAB1, TPSB2, TPSG1, GATA2, MITF, FCER1A*) along the aberrant trajectory, with no induction of key erythroid genes such as *EPOR, KLF1*, and *HES6* relative to MEPs (**Fig. 1H,K**). Collectively these results indicate that loss of CUL3 protein induces fetal hemoglobin in hematopoietic progenitors, but counter-productively, disrupts erythroid commitment. By contrast, reducing CUL3 activity by blocking its neddylation was sufficient to induce fetal hemoglobin while reducing disruption of erythroid commitment in MEPs, as observed with MLN4924.

### Partial inhibition of CUL3 neddylation via DCN1 upregulates fetal hemoglobin without impacting erythropoiesis

Targeting the neddylation pathway could pave the way for new strategies to treat ß-hemoglobinopathies, however, the adverse impact of pan neddylation inhibition on erythroid maturation, dependent on full suppression of CUL3 neddylation, represents a potential undesired risk. We therefore aimed to identify a modulator of the neddylation pathway sufficient to induce fetal hemoglobin while avoiding any adverse impact on erythropoiesis.

To this objective, we integrated a curated neddylation protein-protein interaction network with gene dependency profiles from the DepMap CRISPR viability dataset, reasoning that pathway components whose loss broadly impairs cell viability across diverse cell lines may be less tractable therapeutic targets (*30*). We then pruned nodes whose KO produced substantial viability defects across this dataset, yielding a subnetwork enriched for neddylation regulators predicted to have minimal cytotoxic liability. This subnetwork revealed DCUN1D1, which encodes DCN1 that functions as a scaffold between UBC12 and CUL3 to enhance the transfer of NEDD8 (*31*) (*32*). Because DCN1 functions within the UBE2M-CUL3 axis that we had already implicated in HbF induction, it emerged as a mechanistically prioritized candidate for more selective modulation of this pathway. Consistent with this rationale, DCUN1D1 KO increased HBG mRNA by more than two-fold and increased the fraction of HbF^+^ cells in CD34^+^ cell-derived erythroid cultures (**Fig. 2A,B and fig. S3A,B**).

**Fig. 2.**
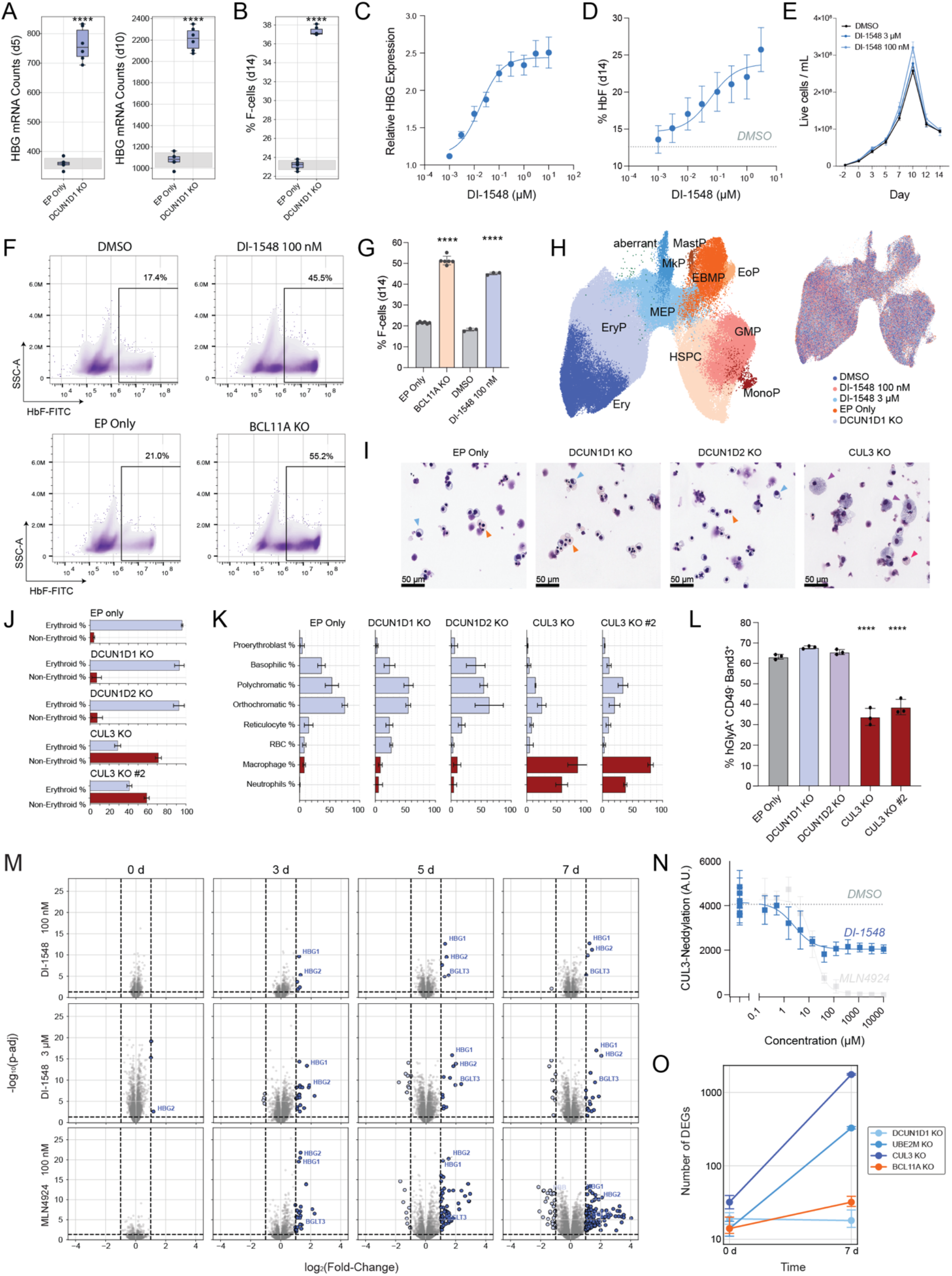
Targeting DCN1 drives adult-to-fetal hemoglobin transition without impacting erythropoiesis. **(A)** Normalized HBG expression measured by nCounter on days 5 and 10. **(B)** % F-cells measured by flow cytometry on day 14. For (A) and (B), statistical significance was determined by unpaired t-test relative to EP Only (****p<0.00005). Grey box denotes 95% confidence interval for the EP Only control. **(C)** Dose-response curve of DI-1548 on HBG expression measured by nCounter. **(D)** Dose-response curve of DI-1548 on % HbF, measured by HPLC. (**E**) Live cell density over the course of differentiation. For (C-E), six biological replicates were collected across four donors. **(F)** Representative flow cytometry gating of cells at day 14, showing HbF-FITC versus SSC-A. **(G)** Quantification of % F-cells by flow cytometry. Statistical significance was determined by unpaired t-test versus experimental control (****p<0.00005). **(H)** UMAP embeddings with cell type annotations from time course scRNA-seq following compound and CRISPR perturbations. **(I)** Bright field images of cytospin slides at day 14. Orange and light blue arrows indicate morphologically normal erythroid intermediate (e.g., orthochromatic and polychromatic erythroblasts); purple and red arrows indicate morphology of macrophage and neutrophil lineage cells, respectively. **(J)** Quantification of erythroid versus non-erythroid cells across cytospin slides. **(K)** Proportion of cells across erythroid maturation stages (light purple) and non-erythroid lineages (red). For (J) and (K), (n=3) (CUL3 KO, n=2). **(L)** Flow cytometry analysis of erythroid maturation markers. Statistical significance determined by ANOVA with Dunnett’s multiple comparisons test (****p<0.00005). **(M)** Volcano plots showing differential gene expression from pseudobulked replicates by timepoint. Dashed lines indicate adj p < 0.05 and |log_2_FC| ≥ 1, respectively. DI-1548 (N=7), MLN4924 (N=3). **(N)** Inhibitory dose response curve of DI-1548 against CUL3 neddylation in HUDEP2 cells, determined by AlphaLISA (N=4). MLN4924 curve underlaid for reference. **(O)** Median number of differentially expressed genes at day 0 and day 7 from scRNA-seq, with bootstrapped 95% confidence intervals (N=3). Panels (A-M, O) were performed in primary human CD34 cells.

DCN1 and its homolog DCN2, coded by DCUN1D2, exhibit strong structural homology, particularly in their UBC12-binding domains and may therefore be targeted in concert by small molecule inhibitors (*33*). We synthesized a previously described inhibitor, DI-1548 (*34*), as an *in vitro* tool compound to test the efficacy of DCN1/2 inhibition in the induction of fetal hemoglobin. Treatment of CD34^+^ cells with DI-1548 exhibited a dose-dependent induction of both HBG mRNA with an Emax of 2.5-fold and an HbF induction of up to 25% with no impact on cell counts (**Fig. 2C-E**).

Furthermore, 100 nM DI-1548 increased the percentage of F-cells to 45.5%, comparable to BCL11A enhancer KO at day 14 (55.2%) with a similar, 2.6-fold increase in either treatment relative to their control (**Fig. 2F,G**). DCUN1D2 KO did not induce fetal hemoglobin, establishing DCN1 as the mechanistic target of DI-1548 for fetal hemoglobin upregulation (**fig. S3C**). Lastly, DCN1 inhibition induced HbF predominantly during the HSPC and/or MEP stage as treating CD34^+^ cells after day 3 of erythropoiesis did not alter fetal hemoglobin (**fig. S3D**).

To interrogate the impact of DCN1 perturbation on various hematopoietic lineage output and erythropoiesis in an unbiased manner, we performed scRNA-seq on perturbed CD34^+^ cells across the same erythroid differentiation time course used in Fig. 1G, sampling days 0, 3, 5, and 7 (**Fig. 2H**). Cell type proportions were highly consistent over time with no aberrant cells (<= 0.001%) detected in either the DCUN1D1 KO or DI-1548 conditions, similar to controls (**fig. S4A,B**). Giemsa staining of CD34^+^ DCUN1D1 KO cultures on day 14 of erythroid differentiation showed cells spanning the full spectrum of erythroblast maturation (blue and orange arrows), with expected stage-specific morphologies and a predominance of erythroid over nonerythroid cells, consistent with preserved erythroid differentiation (**Fig. 2I-K**). Flow cytometry further demonstrated that KO of DCUN1D1 did not impact erythroid maturation (**Fig. 2L**), and a dose response assessment of DI-1548 had no significant impact on the expression of erythroid differentiation markers (*GYPA, TFRC, ITGA4, SLC4A1*) (**fig. S4C**), supporting minimal perturbation of erythropoiesis and skewing to other lineages. We previously noted that the aberrant trajectory formed by CUL3 KO, UBE2M KO, or NAE1 inhibition exhibited induction of myeloid lineage transcription factors (**Fig. 1K**). Consistent with these findings, culturing CUL3 KO CD34^+^ cells in erythroid differentiation media significantly affected cell viability, generated fewer erythroid cells, instead predominantly giving rise to white cells with morphological features of macrophages (purple arrows), neutrophils (red arrows), and other hematopoietic elements (**Fig. 2I-L and fig. S4D,E**).

Comparative differential expression analysis of time-course erythropoiesis using scRNA-seq revealed the striking specificity of DCN1 inhibition in inducing fetal globin expression. At day 7, treatment with 100 nM DI-1548 resulted in significant differential expression of only five genes including HBG1, HBG2, and the globin-regulatory lncRNA BGLT3 compared to the 170 genes differentially affected by 100 nM MLN4924, with similar trends observed at earlier timepoints (adj. p < 0.05 and log_2_(FC) >= 1) (**Fig. 2M**). Given that DCN1 functions as an enhancer of UBC12-mediated neddylation of CUL3, we reasoned that its inhibition would result in more modest reduction in CUL3 neddylation compared to inhibition of NAE1 or UBE2M KO. Indeed, AlphaLISA assessment in HUDEP-2 cells confirmed that DI-1548 partially inhibited CUL3 neddylation, with an E_max_ of ∼ 60% (**Fig. 2N**), and in the erythroleukemia cell line TF1, with an Emax of ∼ 55% (**fig. S5A**). DCUN1D1 KO further confirmed partial inhibition of CUL3, achieving ∼ 57% inhibition (**fig. S5B**). This graded biochemical inhibition was reflected in the corresponding transcriptional impact on erythropoiesis at day 7. CUL3 KO significantly affected the expression of over 1,000 genes, while full inhibition of CUL3 neddylation via MLN-4924 or UBE2M KO altered the expression of hundreds of genes. In contrast, partial inhibition of CUL3 neddylation by DI-1548 or DCUN1D1 KO affected only 5 and 18 genes respectively, similar to the transcriptional specificity of globin switching with BCL11A enhancer knockout (**Fig. 2M,O and fig. S5C**).

NRF2, a key regulator of the antioxidant response, is a well-established CUL3-complex substrate targeted for degradation by the KEAP1 adaptor protein (*35, 36*). Consistent with partial CUL3 inhibition, DI-1548 induced only a modest dose-dependent increase in NRF2 HiBiT signal in a HUDEP2 reporter system, compared to the strong activation observed with Omaveloxolone, an NRF2 activator, or MLN4924 (**fig. S6A**). Although modulation of NRF2 has been implicated in the transcriptional activation of HBG genes (*37*), NRF2 KO only modestly attenuated DI-1548-mediated HBG induction in HUDEP2 cells, suggesting that NRF2 activation alone is insufficient to account for fetal hemoglobin induction following DCN1 inhibition (**fig. S6B**). Together, these results indicate that partial inhibition of CUL3 neddylation modestly stabilizes downstream substrates as observed with NRF2.

### DCN1 inhibition drives locus-specific activation of fetal hemoglobin

As we established that DCN1 inhibition upregulates HBG1 and HBG2 expression, we next characterized chromatin changes at the hemoglobin locus during the early phase of erythroid differentiation. ATAC-seq profiling on differentiation day 5 identified a limited number of sites with significantly increased accessibility, concentrated within the HBG2, HBG1, HBBP1, and BGLT3 regions of chromosome 11 (**Fig. 3A-C, upper panels**). Strikingly, only 0.049% of peaks called genome-wide exhibited a statistically significant change in accessibility after 5 days of differentiation with DCN1 inhibition, with the most prominent changes localized directly to the γ-globin genes and a regulatory region spanning BGLT3 and the HBBP1 transcription start site (|log_2_(FC)| ≥ 1 & adj p < 0.05). This highly selective effect contrasted sharply with MLN4924, which significantly altered 5.9% of peaks called genome-wide at the same thresholds (**Fig. 3A, lower panel and fig. S7**). To assess whether these early accessibility changes were accompanied by acquisition of an active chromatin state and regulatory factor occupancy as differentiation proceeded, we profiled H3K4me3 and NFY-A by CUT&RUN at day 7. H3K4me3 signal increased across the γ-globin genes, consistent with active transcription at this timepoint (**Fig. 3B,C, middle panels**). NFY-A is a known regulator of HBG expression (*38, 39*), and NFY-A enrichment has been linked to gene expression at H3K4me3-marked loci (*40*). Consistent with this model, CUT&RUN profiling at day 7 showed a significantly increased NFY-A binding at HBG2 and HBG1, together with the expected H3K4me3 enrichment pattern across gene bodies (**Fig. 3B,C, lower panels, and fig. S8A,B**). Across all three chromatin profiling studies, the most significant genome-wide changes by both statistical significance and fold change were concentrated at regulatory regions proximal to the γ-globin genes, with few other sites showing significant alterations. Together with our transcriptomics results (**Fig. 2M**), these data demonstrate that DCN1 inhibition induces a highly selective chromatin and transcriptional response centered on the γ-globin locus.

**Fig. 3.**
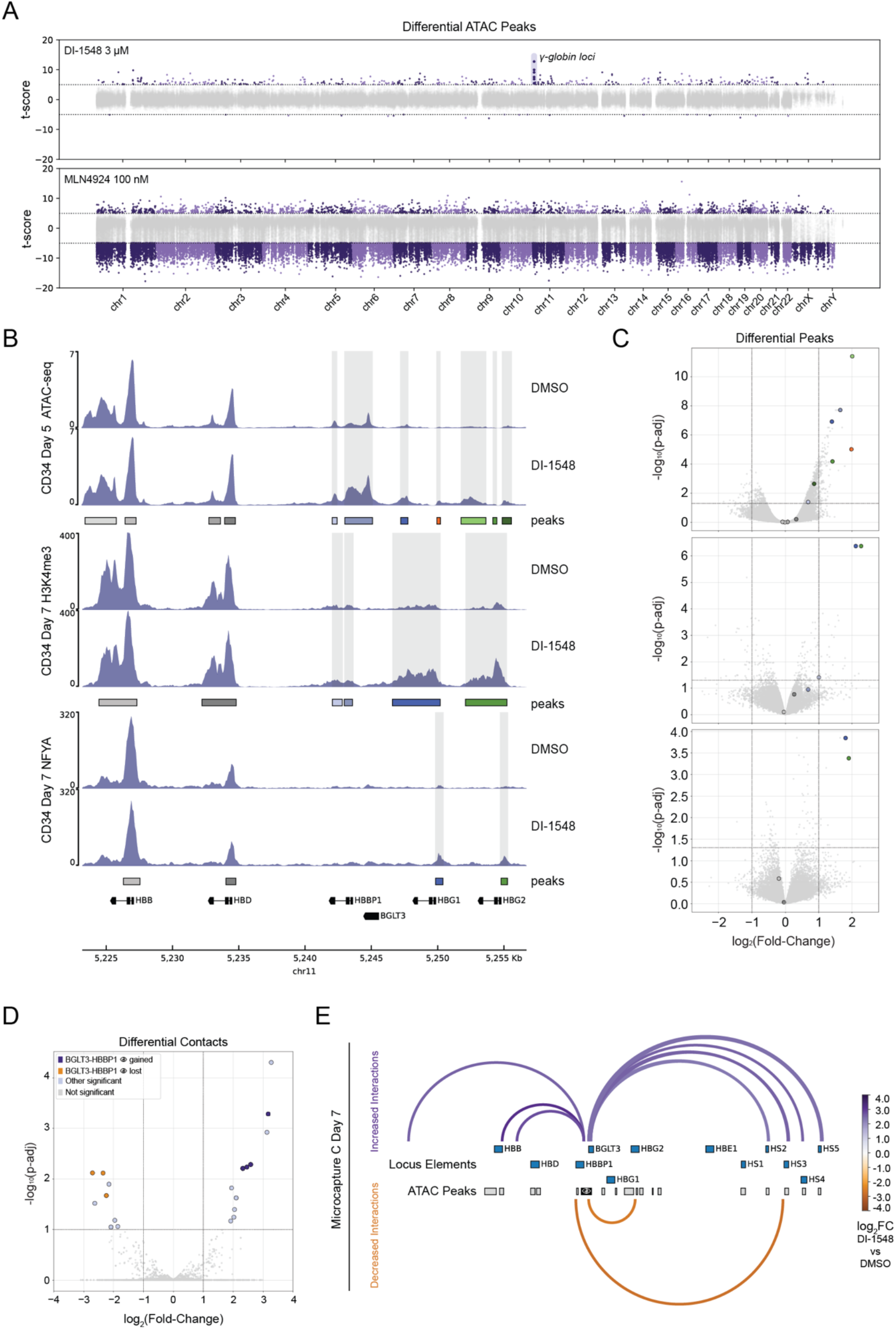
Inhibiting DCN1 induces functional genomic reorganization of the hemoglobin locus. **(A)** Manhattan plot showing t-scores across chromosomes from limma differential peak analysis (N=5). Dotted lines indicate approximate thresholds at ± 5. Peaks spanning HBG2, HBG1, BGLT3, and HBBP1 are enlarged for clarity. **(B)** Track plots of hemoglobin genes showing normalized ATAC-seq and CUTCRUN signal for the indicated antibody and timepoint (ATAC-seq, n=5; CUTCRUN, n=3). **(C)** Volcano plots of genome-wide differential peaks corresponding to the adjacent track plots in (B). Colors of highlighted peaks in (B) match dot colors in (C); all other peaks are shown in grey. (**D**) Differential chromatin contacts inferred from virtual 4C analysis using Micro-Capture-C on day 7 of CD34^+^ erythropoiesis (n=2). **(E)** Significantly altered contacts along the hemoglobin locus and adjacent cis-regulatory regions, visualized from the *BGLT3-HBBP1* viewpoint. 👁 denotes the ATAC-seq peak viewpoint being emphasized in (D), and viewpoint contacts visualized in (E).

Approximately 20kb downstream of the hemoglobin locus exists a cis-regulatory region described as the locus control region (LCR), which consists of five hypersensitive sites (HS1-5) that form measurable and dynamic contacts between the globin locus, as the proteins that link these regions together may be differentially recruited under various contexts (*41*). Micro-Capture-C (MCC) profiling of CD34^+^ cells on day 7 of erythropoiesis, followed by virtual 4C analysis using ATAC-accessible regions as viewpoints, revealed DCN1 inhibition-dependent changes in chromatin contacts at the hemoglobin locus and surrounding cis-regulatory elements (**Fig. 3D and fig. S9A**). Several of the most significantly changed contacts occurred from the BGLT3-HBBP1 viewpoint.

Emerging evidence suggests that the BGLT3-HBBP1 region contains regulatory elements that form and promote chromatin contacts between the LCR and multiple sites across the β-globin locus, facilitating globin transcriptional switching (*42, 43*). Within BGLT3, multiple contacts were significantly gained throughout the LCR HS sites and HBBP1 formed new contacts proximal to HBB (**Fig. 3E and fig. S9B**). Altogether, these genomic studies demonstrate that DCN1 inhibition induces a highly selective reorganization of the globin locus, creating a pro-fetal hemoglobin regulatory conformation with very few other alterations detected genome-wide in complementary assays.

### Superior up-regulation of fetal hemoglobin by combined DCN1 inhibition and HU

Recent advances in SCD therapy have prompted increasing interest in HU-based combinatorial approaches aimed at boosting its efficacy below maximum tolerated dose (*13, 44*). We therefore tested whether DCN1 inhibition could augment HU induced fetal hemoglobin. In cultured human CD34^+^ cell-derived erythroid cells, the combination of DI-1548 (100 nM) and HU (15 μM) induced 26% (mean of n=11) of HbF protein, exceeding the effect of either agent alone (**Fig. 4A, upper panel and B**) and above the threshold of 20% which is considered to be therapeutically meaningful in SCD (*13*). Flow cytometric analysis showed a higher proportion of HbF^+^ cells (69.5%; mean of n=3) in the combination-treated group (**Fig. 4A, lower panel**). Notably, the combination also induced a rightward shift in intracellular HbF intensity, consistent with pan-cellular HbF expression. These findings demonstrate DCN1 inhibition can significantly enhance HbF induction by HU.

**Fig. 4.**
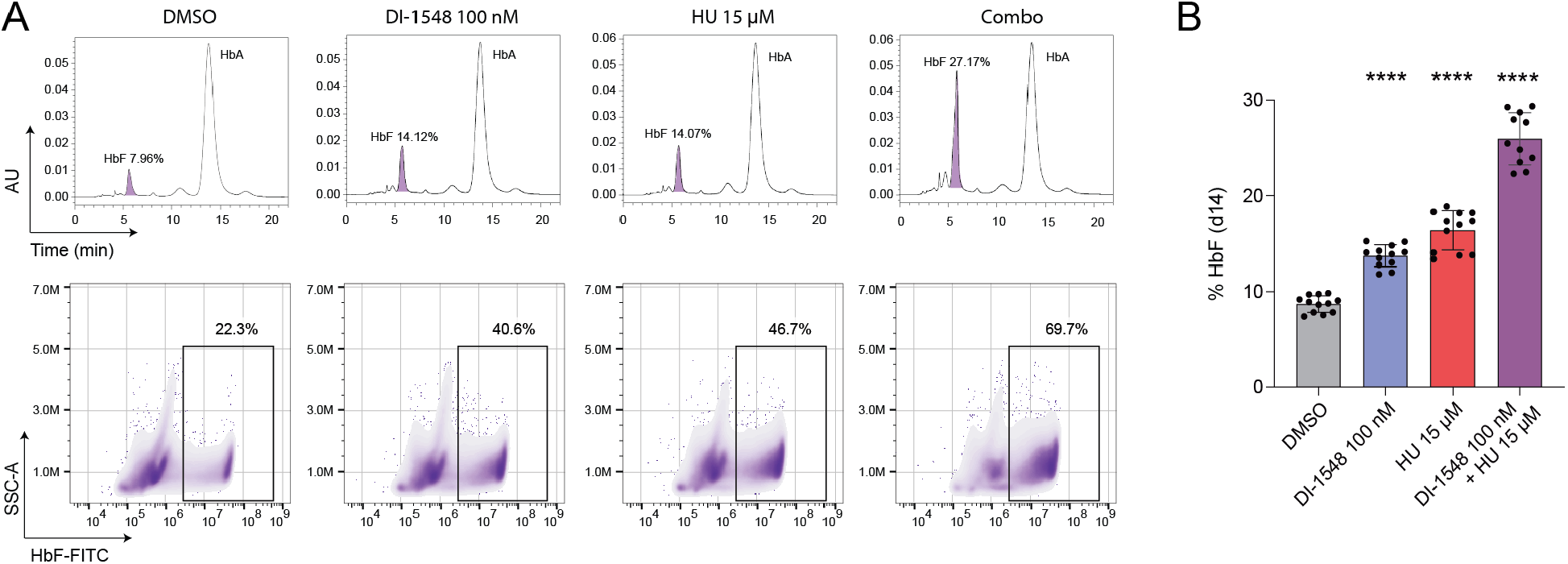
Inhibiting DCN1 works cooperatively with HU in the induction of fetal hemoglobin. **(A)** Top: Representative HPLC chromatograms from day 14 cultured erythroid cells showing fetal hemoglobin (HbF) and adult hemoglobin (HbA). Bottom: Representative flow cytometry analysis of intracellular HbF in day 14 erythroid cells (n=3). **(B)** Percentage of HbF quantified by HPLC in day 14 erythroid cells treated under indicated conditions. Data represent biological replicates from four independent studies using cells from a single donor. Statistical significance was determined using one-way ANOVA with Dunnett’s multiple comparisons test (****p<0.00005).

### Novel DCN1 inhibitor CLY-124 induces fetal hemoglobin and synergizes with HU in a humanized mouse model

DI-1548 is an *in vitro* tool compound for cell-based studies; however, its utility for *in vivo* assessments was impeded by its reported rapid clearance (*34*). Integrated medicinal chemistry efforts aided by structure-based design informed by X-ray crystallography combined with comprehensive analytical and cell-based characterization culminated in the discovery of CLY-124, a potent covalent inhibitor of DCN1 with optimized ADME and pharmacokinetic properties (Fig. 5A and Table S1). X-ray crystallography demonstrated CLY-124 binding to the purified C-terminal Potentiating Neddylation (PONY) domain (*45*), the site that interacts with UBC12 N-terminal 12-mer peptide associated with transfer of NEDD8 to Cullins. The co-crystal structure of CLY-124 in complex with DCN1 was resolved to 2.48 Å, and the electron density of the bond between the sulfur of Cys115 and the C of the nitrile confirmed covalency. The specificity of CLY-124 binding to DCN1 was further confirmed using LC-MS analysis revealing both unmodified DCN1protein (23124.91 Da) and the CLY-124 – DCN1 adduct (23714.65 Da) (**fig. S10A**).

**Fig. 5.**
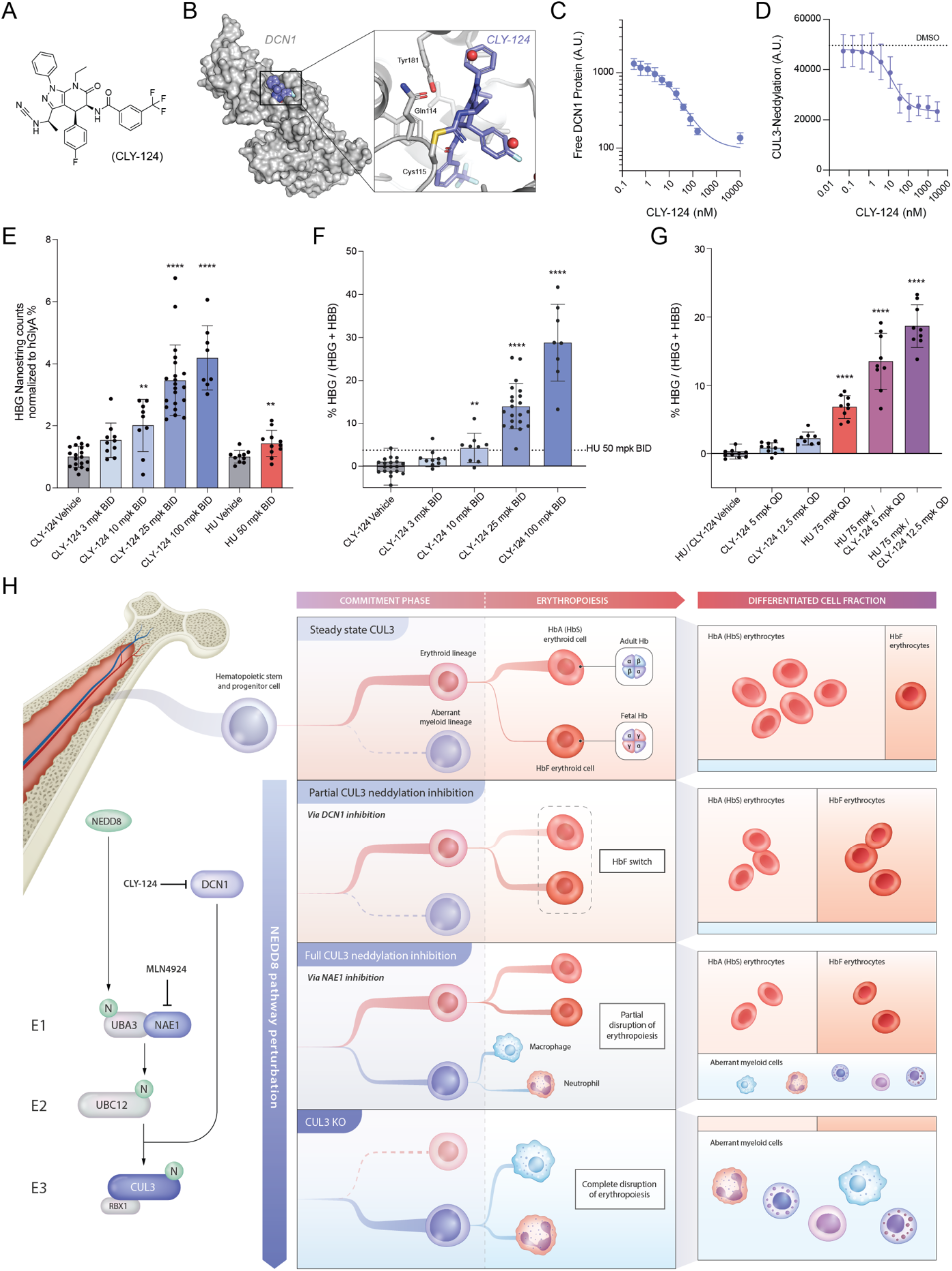
CLY-124 is a potent, selective, and irreversible covalent inhibitor of DCN1 for in vivo HbF induction. **(A)** Chemical structure of CLY-124. **(B)** Co-crystal structure of CLY-124 bound to DCN1. **(C)** Dose-dependent increase in DCN1 occupancy by CLY-124 in TF-1 cells, measured by AlphaLISA signal loss (n=3). **(D)** Dose-dependent reduction in Cul3 neddylation by CLY-124, assessed by AlphaLISA signal with DMSO set at 0 %. Data represent mean ± SD (n=27). **(E)** HBG gene expression analysis. **(F)** Globin ratio expressed as HBG transcripts as a percentage total B-globin (HBG+HBB). Data in (E) and (F) were combined from two studies: Study 1 tested 25 and 100 mpk CLY-124 and 50 mpk hydroxurea (HU); Study 2 tested CLY-124 at 3, 10, and 24 mpk. All doses were administered BID. **(G)** Globin ratio (HBG / (HBG + HBB)) in NBSGW mice treated with CLY-124 (5 or 12.5 mpk), HU (75 mpk), or combinations thereof. (F-G) Data obtained by nCounter gene expression analysis from bone marrow of NBSGW mice engrafted with human CD34^+^ cells and treated for 22 days starting 12-13 weeks post-engraftment. For (E-G) statistical significance was determined using unpaired t-test or one-way ANOVA with Dunnett’s multiple comparisons test, as appropriate (**p<0.005, ***p<0.0005, ****p<0.00005). **(H)** Schematic illustrating the effects of perturbing distinct nodes of the neddylation pathway on erythropoiesis and fetal hemoglobin induction.

Potency of CLY-124 binding to DCN1 was measured using a competitive TR-FRET with a non-covalent DCN1 binder, DI-591 (*33*) with an IC50 of ∼80 nM at 1 hour and ∼4.98 nM at 24 hours of incubation (**fig. S10B**). Due to the time dependent shift in the IC50, the covalent interaction was further characterized by measuring k_inact_/Ki (*46*). In a time-course TR-FRET assay, CLY-124 interacted with DCN1 with a k_inact_/Ki of ∼0.004647 nM-1*min-1 (**fig. S10C and Table S1**). Similarly, CLY-124 interacted with DCN2 with a k_inact_/Ki of 0.0025 nM-1*min-1 (**fig. S10D and Table S1**). Selective binding of CLY-124 to all DCN isoforms was tested by Surface Plasmon Resonance (SPR), which demonstrated association to DCN1 (KD = 13.9 nM) and to DCN2 (KD = 41.5 µM) and a complete lack of binding to DCNs 3, 4, and 5. (**Table S1 and fig. S11A-G**). In TF-1 cells, CLY-124 occupied DCN1 in a dose-dependent manner with an IC50 of 10.15 nM (**Fig. 5C**) and led to a concomitant inhibition of CUL3 neddylation, with an IC50 of 15 nM and an Emax of 60% up to a tested concentration of 10 µM suggesting a partial disruption of this process (**Fig. 5D**). In summary, these results highlight the structural and functional basis of DCN1 inhibition by CLY-124.

The NBSGW mouse model, which captures engrafted human hematopoiesis in the murine bone marrow (*47*), was utilized to investigate the impact of DCN1 inhibition on human fetal hemoglobin induction *in vivo* (**fig. S12A**). To this end, mobilized peripheral blood CD34^+^ cells from healthy volunteers were intravenously infused to establish a human xenograft in 8-week-old NBSGW mice. After 12 - 13 weeks, mice were randomized based on % hCD45 engraftment levels and subjected to oral administration of either CLY-124 or HU at the indicated doses for 21 days. CLY-124 was well tolerated and dosed mice showed body weights comparable to vehicle treated mice throughout the study (**Fig S12B, C**). NBSGW mice in all the *in vivo* studies showed comparable levels of hCD45 engraftment in the bone marrow regardless of compound or vehicle treatment (**fig. S12D and G**). *In vivo* exposure of CLY-124 measured from a subset of 25 mg/kg (mpk) twice a day (BID) and 100 mpk BID groups demonstrated dose-dependent increase in plasma compound levels (**fig. S12E**).

CLY-124 elicited a dose dependent induction of HBG gene expression in human CD34^+^ cell-derived erythroid cells within the bone marrow (**Fig. 5E**). HBG gene expression ranged from 2-fold in the 10 mg/kg BID treated mice to 4-fold in both 25 mg/kg and 100 mg/kg BID treated mice (**Fig. 5E**). In comparison, HU at 50 mg/kg BID demonstrated a 1.4-fold induction in HBG gene expression (**Fig. 5E**). Administration of CLY-124 did not suppress erythropoiesis in comparison to HU, which significantly lowered Ter119^+^ murine erythroblasts in the bone marrow (**fig. S12F**), recapitulating the unperturbed erythropoietic differentiation in the presence of DCN1 inhibition *in vitro* (**Fig 2L**). Notably, CLY-124 increased the HBG-to-total globin ratio, with the highest tested dose of 100 mg/kg BID achieving a mean of 28.8% (**Fig. 5F**). This magnitude of induction, measured as the HBG-to-total globin ratio in the humanized NBSGW model, is comparable to that reported for recent therapeutic approaches to SCD, including BCL11A +58 enhancer targeting (*48*).

To examine if the *in vitro* cooperativity with HU translates, humanized NBSGW mice were co-treated in a staggered treatment regimen (Methods) with CLY-124 and HU, with CLY-124 doses derived from the lower end of the dose response tested in the previous study (**Fig. 5 E, F**). Combination treatment with HU with low doses of CLY-124 at 5 mpk and 12.5 mpk once-a-day (QD), significantly increased the percentage of HBG over total β-globin transcripts and was superior to either agent alone compared to vehicle (**Fig. 5G**). Two-way ANOVA with interaction terms revealed statistically significant synergistic effects between HU 75 mg/kg and CLY-124 at both tested doses. Specifically, the interaction between HU 75 mg/kg and CLY-124 5mg/kg was significant (interaction estimate = 5.87; 95% CI: 2.93-8.81; p = 2.04 × 10^-4^), and the interaction between HU 75 mg/kg and CLY-124 12.5 mg/kg was even stronger (interaction estimate = 9.62; 95% CI: 6.59-12.64; p = 5.84 × 10^-8^), indicating dose-dependent synergy between HU and CLY-124. CLY-124 as a monotherapy was a more potent inducer of fetal hemoglobin than HU and when combined with HU, synergistically increased the HBG gene ratio in vivo. Collectively, these findings support a model in which profound loss of CUL3 activity disrupts steady-state erythropoiesis, whereas partial loss of CUL3 activity through DCN1 inhibition induces HbF without detectable impairment of erythropoiesis (**Figure 5H**).

## DISCUSSION

Beta-hemoglobinopathies namely thalassemia and sickle cell disease are major red blood cell disorders affecting several millions of patients, with high unmet need. Sickle cell disease (SCD) was one of the earliest molecular diseases described in medicine, with the first clinical report published in 1910 (*49*). More than a century later, it affects over 7.5 million individuals worldwide and continues to impose a major burden of early mortality, particularly in children under 5 years of age (*50*). Notably, higher HbF levels are associated with reduced childhood mortality and increased life expectancy in patients with SCD (*5*).

Here, we characterized an unappreciated role for a Neddylation axis involving NEDD8, UBC12, CUL3, and DCN1 in fetal hemoglobin regulation. We identified DCN1 as a therapeutically viable target to modulate the Neddylation axis and designed and optimized a first-in-class DCN1 inhibitor, CLY124. In contrast to pan-neddylation inhibitors, DCN1 inhibition either genetic or pharmacological, resulted in partial suppression of CUL3 neddylation. This led to a dose-dependent induction of HbF through specific and selective regulation of the HBG1/2 loci involving lncRNAs, BGLT3/HBBP1, and known activators, namely NFYA, through differential contacts between the locus chromatin elements with the upstream LCR. This regulatory mechanism culminated in a remarkably selective induction of HBG1/2, with a narrow transcriptional footprint comparable to the BCL11A +58 enhancer KO. This stands in contrast to other recently reported HbF-inducing interventions such as WIZ inhibition (*51*), EED knockout (*52*), or treatment with HU (*53*), as well as broad spectrum neddylation inhibition using MLN4924 and CUL3/UBE2M KO, which results in widespread transcriptomic changes, reflecting their broad systemic impact. DCN1 inhibition had no significant impact on *in vitro* erythroid differentiation or murine erythropoiesis, consistent with a prior report showing no bone marrow pathology in transgenic DCN1 knockout mice (*54*). Finally, we demonstrate that CLY-124 induces HbF in the humanized NBGSW mouse model superior to HU as monotherapy and in synergy with HU as a combo.

The role of neddylation in HbF regulation, and DCN1 as a target for HbF induction, has not previously been described. However, individual components of the CUL3 neddylation axis have already been linked to HbF control. Consistent with our findings in primary CD34^+^ erythroid cells, prior work in HUDEP-2 cells showed that disruption of CUL3 increases HbF and specifically implicated the BTB adaptor SPOP in this effect (*55*). More broadly, CUL3 functions as a scaffold for a large family of BTB-domain substrate adaptors that recruit distinct proteins for ubiquitination and proteasomal degradation (*56*). Among these pathways, both NRF2 and SPOP have been implicated in regulation of HBG expression in erythroid cells (*57*) (*55*). Since DCN1 inhibition produced only modest NRF2 activation, likely through reduced CUL3 activity toward KEAP1-dependent NRF2 turnover, we hypothesize that HBG induction by DCN1 inhibition reflects partial attenuation of multiple CUL3 adaptor pathways rather than NRF2 stabilization alone. These may include KEAP1 and SPOP, as well as additional, as-yet-unidentified CUL3-dependent mechanisms that remain to be defined.

Pan-neddylation inhibition with the NEDD8-activating enzyme (NAE) inhibitor MLN4924 up-regulated HbF at concentrations as low as 10 nM. However, at 100 nM and above it caused marked cytotoxicity and impaired erythroid maturation, leaving only a narrow therapeutic window. Such on-target toxicity mirrors clinical experience with NAE inhibitors, in which anemia was one of the most frequent adverse events (*58, 59*). Mechanistically, our data indicate that this liability results from broad suppression of CUL3 activity as genetic CUL3 KO produced the most profound defects in erythroid differentiation and cell viability. In contrast, pharmacologic or genetic inhibition of DCN1 in primary erythroid cells reduced CUL3 neddylation by no more than ∼60%, yet this partial inhibition was sufficient to increase HbF expression without any detectable impairment of erythropoiesis. We hypothesize that this restrained level of CUL3 inhibition allows stabilization of specific CRL3 substrates that promote HbF while maintaining enough cullin-mediated proteostasis to support red-cell maturation. More broadly, our findings suggest that targeting a regulatory co-factor such as DCN1 can serve as a biologically constrained way to modulate a more toxic target; by attenuating rather than abolishing CUL3 activity efficacy was achieved without the dose-limiting toxicities seen with pan-neddylation inhibition. This indirect targeting strategy may be generalizable to other therapeutic pathways where directed inhibition is limited by narrow safety margins.

Our chromatin profiling mapping NFYA binding, revealed that DCN1-dependent regulation of the HBG1/2 loci involves changes in NFYA occupancy at the y-globin promoter. NF-Y is a heterotrimeric transcription factor that binds CCAT motifs, and its DNA-binding specificity is conferred by the NFYA subunit (*60*). Binding of NF-Y/NFYA to the proximal CCAAT box of the y-globin promoter is known to activate HBG transcription (*61*). This activator is antagonized by HbF repressors such as BCL11A and ZBTB7A (also called LRF), which bind to adjacent motifs (-115 and -200, respectively) and recruit corepressors to displace NF-Y and silence HBG1/2 (*39*). Depletion of BCL11A allows

NF-Y to reoccupy the promoter (*39*), highlighting the dynamic competition between NFYA and these repressors. Partial inhibition of cullin neddylation via DCN1 could therefore shift the balance by altering the levels or activity of the repressors. Future studies should test how DCN1 inhibition affects the stability and post-translational modifications of these repressors and whether this contributes to fetal hemoglobin induction.

*In vivo* studies with NBGSW mice harboring human CD34^+^ cells, demonstrated dose dependent induction of HBG superior to HU. In our studies, low doses of DCN1 inhibitors both *in vitro* and *in vivo*, augmented the HbF induction response induced by HU. Combination with a DCN1 inhibitor could potentially lower dosage of HU in patients, thus mitigating some of HU’s side effects while maintaining a superior HbF induction profile. Collectively, these findings support DCN1 and the neddylation pathway more broadly as a specific regulator of globin switching and a potential therapeutic target in hemoglobinopathies especially SCD, capable of inducing selective genomic reorganization at the globin locus to promote fetal hemoglobin expression. In combination with a supportive preclinical toxicology dataset, these data have inspired further clinical development of CLY-124 as a promising compound for induction of HbF. CLY-124 has now entered a first-in-human study evaluating safety, pharmacokinetics and varied assessments of HbF in healthy volunteers and participants with SCD.

## Acknowledgements

We thank past and current members of Cellarity who have contributed to the discovery of DCN1 and the development of CLY-124. Additionally, we thank Dr. Alex Shalek, Dr. Leonard Zon, and Dr. Vijay Sankaran for their thoughtful review of this manuscript.

## Author Contributions

Conceptualization: SAM, SK, RN, KM, AT, MC, and CT

Methodology: SAM, AJM, EHZ, SL, OL, SM, CH, RN, CD, AM, KM, NL, RU, AS, SK, MR, CH, JJO, XS, FL, QT, YY, WZ, TF, and MC

Visualization: SAM, SK, and KM

Project administration: SAM, SK, AJM, EHZ, SL, RN, JJO, AT, and MC

Supervision: SAM, SK, AJM, EHZ, SL, JJO, XS, AT, MC, CT

Writing-original draft: SAM and SK

Writing-review & editing: SAM, SK, AT, MC, and CT

## Data, Code, and Materials Availability

Structural data will be deposited in the RCSB Protein Data Bank. Raw ATAC-seq, CUT&RUN, and Micro-Capture C data will be deposited in the Sequence Read Archive. Mapped count matrices and intermediary files will be made available through the Gene Expression Omnibus. A processed and integrated scRNA-seq data object will be made available through the Broad Single Cell Portal.

## List of Supplementary Materials

Materials and Methods

Figs. S1 to S12

Tables S1 to S4

References *(1-22)*

