## Supplemental Materials and Methods for "DCN1 inhibitor induces fetal hemoglobin through self-limited regulation of CUL3 neddylation"

**Cells and Erythroid Differentiation**

Hematopoietic stem and progenitor cells (HSPCs) were obtained from peripheral blood leukopaks mobilized with granulocyte colony-stimulating factor (G-CSF), plerixafor, or G-CSF and plerixafor (Combo) from StemCell Technologies (Vancouver, BC, Canada). $CD34^+$ cells were enriched from leukopaks by positive immunomagnetic separation and cryopreserved in serum free medium. $CD34^+$ cells were cultured using a 3-phase erythroid culture system with routine media changes as essentially outlined in (*1*) which consisted of $CD34^+$ progenitor cell expansion, erythroid expansion, and differentiation. Compound incubation studies were carried out by adding compounds to the cells after 2 days of thawing and while in the $CD34^+$ progenitor cell state. Compounds were replenished in all the studies every 2 or 3 days whenever there is a medium change and were maintained in cell culture medium until study termination. TF-1 cells were purchased from ATCC (Cat No. CRL-2003) and cultured in RPMI-1640 cell culture medium containing 10 % FBS and antibiotics. HUDEP-2 cells obtained from the Riken Institute were cultured based on published methods (*2*).

**Cul3 Neddylation AlphaLISA assay**

AlphaLISA assay for detecting CUL3 neddylation was performed according to manufacturer's instructions (Revvity, Hopkinton, MA). Briefly, approx. 17K TF1 cells or 20 K HUDEP-2 cells were plated per well of a 384-well plate in Iscove's Modified Dulbecco's Medium (IMDM) without supplements and were treated with compounds at a range of concentrations from 0.05 nM to 3,000 nM at 3-fold dilutions in DMSO for 3 hours. Cells were subjected to lysis with 5 X AlphaLISA lysis buffer (Revvity, Hopkinton, MA). CUL3 neddylation was detected by addition of biotinylated anti-NEDD8 antibody followed by AlphaLISA acceptor beads conjugated with anti-CUL3 antibody following manufacturer's protocols (Revvity, Cat No. CUSB236139000EA). After overnight incubation, Streptavidin-coated Alpha Donor beads (Revvity cat no. 6760002S) were added, and the plates were read on a VICTOR Nivo Multimode Microplate Reader (Revvity, Hopkinton, MA). The dose-response curves were fitted with the Hill equation to obtain IC50 values using Graphpad Prism software.

**Gene Expression Analysis using nCounter technology by Nanostring®**

Mobilized peripheral blood CD34 cells from healthy human donors were differentiated into the erythroid lineage, with compounds added after 2 days of culture and replenished every 48 – 72 hours. Compound incubations were performed at concentrations ranging from 0.003 uM to 10 uM for 12 days, after which cells were harvested. On the terminal cell harvest day 100K cells were collected and lysed in 25 µL of RLT (QIAGEN, Cat No. 79216) with 1x β-Mercaptoethanol (Gibco, Cat No. 21985-023) and placed on a plate shaker at 300-500 RPM for 5 minutes at room temperature. Similarly, bone marrow cell pellets (total 500K cells) from NBSGW mice were lysed in 25 µL of RLT (QIAGEN, 79216) with 1x β-Mercaptoethanol (Gibco, 21985-023). Cell lysates from both in vitro and in vivo samples were then tested for gene expression using the nCounter technology by Nanostring, following manufacturer's instructions. All hybridizations were done in a total volume of 15 µL (3 µL of RNA lysate added to master mix of 12 µL probe A/B, capture probe /reporter probes, proteinase K and attenuation oligos suspended in hybridization buffer). Expression of genes tested include human HBG1, HBB, GYPA, TFRC, ITGA4, and SLC4A1 which were normalized to housekeeping genes ABCF1, GUSB, POLR2A, RPL19, and SDHA. The probe sequences for each of these genes from the nanostring design is provided in the table below (Supplementary Table 2).

**HPLC based measurement of HbF protein**
Erythroid cell pellets from in vitro cultures were collected on day 18 of erythroid differentiation with compound incubations. From in vivo experiments, bone marrow erythroid cells were enriched using immunomagnetic separation for human GLYA$^{+}$ (CD235a) cells. In both cases, 500 K cells were collected, centrifuged at 300 g for 5 mins, and supernatant aspirated and removed. Samples were then injected into the HPLC system. Hemoglobin separation was done using Cation exchange HPLC with a Synchropak CM-300, 250 mm x 4.6 mm, 6u column (Promigen Life Sciences, LLC) containing a negatively charged resin, which binds to the positively charged hemoglobin molecules. The mobile phase, which is a buffered salt solution, was adjusted to control the ionic strength and pH, facilitating optimal separation. Separated hemoglobin peaks were detected by UV absorbance at wavelengths specific to each of the hemoglobin species (415 nm), and the species names assigned by the software which was included as output.

**Flow Cytometry analysis for cell surface markers and intracellular HbF**
*Flow cytometry analysis of erythroid differentiation.* Following 14 days of erythroid differentiation, cells were analyzed for cell surface expression of maturation markers by resuspending them with an antibody cocktail consisting of anti-CD71-PE (BD Bioscience, Cat. No 566722, clone OKT9), anti-CD235a(GlyA)-PE-Cy7 (BioLegend, Cat No. 306620, clone HIR2), anti-CD49d-BV421 (BD Bioscience, Cat No. 565277, clone 9F10), anti-CD233 (Band3)-FITC (IBGRL, Cat No. 9439, BRIC6) and anti-CD45-PerCP (BD Bioscience, Cat No. 347464, clone 2D1) by flow cytometry (Supplementary Table 3). Cells were centrifuged with PBS for 5 min at 300 g, followed by incubation with the antibodies diluted in PBS+ 0.5% BSA + 2mM EDTA for 25 min at 4 °C in the dark. Excess of antibody was centrifuged with excess of PBS and resuspended in 200µL of PBS+ 0.5% BSA + 2mM EDTA. All samples were acquired using a Novocyte Quanton instrument (Agilent) and analyzed with FlowJo (Becton, Dickinson & Company).

*Measuring Intracellular levels of HbF.* ~$1x10^{6}$ bone marrow cells were placed in a 96-well U-bottom plate and centrifuged with PBS+ 0.5% BSA + 2mM EDTA for 5 min at 300 g, followed by fixation with 200 µL of 0.05% freshly diluted glutaraldehyde (Sigma-Aldrich, G5882) in PBS for 10 min at room temperature (RT). After fixation, the cells were centrifuged 2 times with PBS + 0.5% BSA + 2mM EDTA. Cells were stained for human erythroid cell surface markers using 5 µL anti-CD71-PE (BD Bioscience, #566722) and 5 µL of anti-CD235a-PeCy7 (BioLegend, Cat No. 306620, clone HIR2) in 40 µL of PBS + 0.5% BSA + 2mM EDTA. Cells were incubated for 15 min at 4 °C in the dark. After centrifuging the cells with 200 µL of PBS + 0.5% BSA + 2mM EDTA, cells were permeabilized by resuspending in 200 µL of 0.1% Triton X-100 (Life Technologies, HFH10) solution in PBS + 0.5% BSA + 2mM EDTA for 5 min at RT and then centrifuged with 200 µL of PBS + 0.5% BSA + 2mM EDTA. HbF intracellular staining was performed with 10µL anti-HbF-FITC (Life Technologies, MHFM01-4) antibody in 100µL PBS + 0.5% BSA + 2mM EDTA for 15 min at RT then centrifuged with 200µL PBS + 0.5% BSA + 2mM EDTA and resuspended in 200µL PBS + 0.5% BSA + 2mM EDTA. All samples were acquired using a Novocyte Quanton instrument (Agilent) and analyzed with FlowJo (Becton, Dickinson & Company).

**CRISPR-based gene editing**
*Gene knockouts.* CD34 cells were subjected to CRISPR based editing 48 hours after thawing as outlined in fig. S1A. Briefly, ribonucleoprotein (RNP) complexes were prepared by incubating Cas9 nuclease (GenScript) and guide RNAs (Synthego or GenScript) at a ratio of either 1:1 or 1:2. Sequences of gRNAs for various genes used in this study are listed in (Supplementary Table 4). Cell

aliquots (50 K) on day -2 of erythroid differentiation were resuspended in 20 ml of P3 reaction buffer and combined with 5 µl of RNP mixture. Cell – RNP mixture was then mixed and added to the electroporation cuvette and was subjected to electroporation in a 4D nucleofector (Lonza #AAF-1002B), using the program CA137. Post-electroporation, 100 µl of complete medium was added to the cuvette and cells transferred to a tissue culture 96-well plate pre-loaded with 100 µl of warm medium. Aliquots of cells at 50 – 100 K were collected on days 0, 3, 5, 7, 10, and 12. Genomic DNA from these cells were isolated using gDNA isolation kit (Lucigen) and subjected to PCR to detect knockout efficiency. NFE2L2 KO in HUDEP2 cells was performed using a single CRISPR guide per manufacturer's instructions (Synthego), and knockout was validated by genomic sequencing with indel percentages above 90% in four consecutive cell passages.

*Endogenous HiBiT tagging to NRF2 in HUDEP2 cell line.* 50x10^3 viable HUDEP2 cells were transfected with RNP complex [1.16 µM Hifi Cas9 and 3.3 µM gRNA(Hs.HC9.BMRS1779.AA ) targeting NRF2], 4 µM ssODN(CD.HC9.FKBK3937) and 4 µM Alt-R Cas9 Electroporation enhancer by electroporation, followed by culturing in expansion media with 1.7µl /ml Alt-R HDR Enhancer V2. Medium exchange was performed overnight post- electroporation without Alt-R HDR Enhancer. Successful integration of the HiBiT tag was determined 48 h after editing by measuring luminescence in cell lysates in the presence of LgBiT protein followed Promega manufacture instruction (Nano-Glo® HiBiT Lytic Detection System). To determine the effect of compounds on NRF2, HUDEP2 NRF2-HiBiT cells were plated in alphaPlate-384 shallow well (Revvity 6008350) at 20,000 cells per well in IMDM (GIBCO, 12440-053) with 1% fetal bovine serum. MLN4924, omaveloxolone and DI-1548 were serially diluted in DMSO and delivered to cells at 0.17 to 10,000 nM, along with vehicle control; final DMSO concentration was 0.1%. After overnight incubation at 37 °C and 5% $CO_2$, HiBiT detection reagents were mixed and delivered to the culture cells per manufacturer instruction (Nano-Glo® HiBiT Lytic Detection System, N3030, Revvity). The plate was shaked for 10 minutes at room temperature and the luminescent signal was read on Victor Nivo (Revvity).

**Western blotting analysis**

Western blotting analysis for protein expression was performed from cell pellets ranging from 0.5 – 1 M cells derived from erythroid cultures of human CD34 cells. Briefly, cell pellets were centrifuged and rinsed with PBS to remove residual culture medium and lysed with RIPA cell lysis buffer (Cell Signaling #9806) containing PMSF and SDS. Protein levels in cell lysates were measured using BCA protein assay. Protein samples were resolved using a precast 10% Criterion Bis-Tris SDS-PAGE gel (BioRad #3450113) and separated proteins were transferred to Immobilon® PVDF membranes (Millipore, # IPFL00010). Proteins were probed with specific antibodies against CUL3 (Cell Signaling #10450), CUL1 (Cell Signaling #4995), DCN1 (Abcam #ab181233), and UBC12 (Santa Cruz Biotechnology #sc-390064) normalized to GAPDH (Cell Signaling #2218) and visualized using an Odyssey® F instrument (Licor).

**LC/MS analysis of protein binding**

Covalent binding of CLY-124 to purified DCN1 protein was tested using LC/MS. Recombinant DCN1 protein was expressed in E. coli with a His-tag and purified using a Ni-NTA column. The tag was then cleaved using a His-tag TEV protease and removed using a second Ni-NTA column. Protein purity was verified with SDS-PAGE and intact MS. Purified recombinant DCN1 protein was diluted in a buffer containing 25 mM Tris-HCl, 200 mM NaCl, and 1 mM DTT at a concentration of 400 nM. To this solution, 0.4 µL of CLY-124 in DMSO was added with a final DMSO concentration of

1 %. CLY-124 binding to DCN1 was tested with a 11-point dose response with 3-fold serial dilution starting at a top concentration of 10 μM. The reaction was performed for 3 h in the dark at room temperature and stopped after 3 h with formic acid at a final concentration of 0.2 %. Quenched assay plates were analyzed with an Agilent RapidFire 360 system connected to an Agilent 6545 Q-TOF mass spectrometer equipped with an AJS source. 10 mL of sample volume was loaded onto a custom packed cartridge (4 mL, PLRP-S 30 mm/1000 Å pore; Optimize Technologies) with loading buffer (dd$H_2O$ with 0.09 % (vol/vol) formic acid and 0.01 % (vol/vol) trifluoroacetic acid; 1.25 ml/min) for 6 seconds before being eluted directly into the mass spectrometer in elution buffer (80% acetonitrile with 0.09% (vol/vol) formic acid and 0.01% (vol/vol) trifluoroacetic acid; 0.5 ml/min) for 7 s. The cartridge was re-equilibrated with loading buffer for 1 s before collection of the subsequent sample. The Q-TOF was operated in TOF-only positive ionization mode set to the following parameters: Gas Temp = 350 °C, Drying Gas = 7 l/min, Nebulizer = 50 psi, Sheath Gas Temp = 400 °C, Sheath Gas Flow = 12 l/min, VCap = 4000 V, Nozzle Voltage = 1000 V, Fragmentor = 125 V, Skimmer = 65 V and Oct 1 RF Vpp = 750 V. Raw MS data files were deconvoluted and analyzed using the Agilent MassHunter Bioconfirm software package (Bioconfirm version 9) to identify both parent protein and expected compound adduct mass signatures. MS heights of the parent protein and the adduct were designated as P and A, respectively. The percentage of adduct formed was then calculated as A / (P+A).

**SPR for analysis of DCN interaction**

SPR experiments performed to investigate binding of CLY-124 to the 5 isoforms of DCN namely DCN1, 2, 3, 4, and 5 on a 8K Biacore instrument (Catalog:29722782, Cytiva). Biotinylated DCN recombinant proteins were immobilized on Series S Sensor Chip SA(BR100531) with running buffer containing 20 mM Tris-HCl, pH 7.5, 150 mM NaCl, 1 mM DTT, 0.005 % BSA, 0.05 % Tween20. The amine-PEG-biotin was dissolved in HBS-N (10 mM HEPES pH 7.4, 150 mM NaCl) or HBS-P+ (10 mM HEPES pH 7.4, 150 mM NaCl, 0.05% Tween20). Compounds were serially diluted 2-fold in the running buffer with final DMSO concentration of 2 % and injected into the flow cells on a flow rate of 30 μl/min for 90 s (association), followed by running buffer for 360 s dissociation time. KD values were obtained with 1:1 binding mode using Biacore™ Insight Software.

**DCN1 target occupancy assessment using competition AlphaLISA**

We developed a competition-based AlphaLISA assay to study DCN1 target occupancy. In brief, recombinant anti-DCN1 antibody (Abcam, #ab250448) was conjugated to unconjugated AlphaLISA acceptor beads per manufacturer instructions (Revvity, #6772002). A second DCN1 antibody was biotinylated (Antibodies-online, #ABIN5608786). After incubation with DCN1 inhibitors for 3 hours at 37 °C and 5% CO2, TF1 cells were lysed with AlphaLISA lysis buffer (Revvity, #AL003F). Anti-DCN1 acceptor beads and biotinylated anti-DCN1 antibody were added to cell lysates, both diluted in 1X Immunoassay buffer (Revvity, #AL000F). This mixture was incubated at room temperature overnight before the addition of AlphaScreen Streptavidin-coated Donor beads (Revvity, #6760002) diluted in 1X Immunoassay buffer. After 1-hour incubation at room temperature, plates were read out using Victor Nivo plate reader (Revvity).

**TR-FRET based assessment of CLY-124 interaction with DCN1: IC50 and Kinact/KI Measurements**

TR-FRET based determination for CLY-124 interaction with DCN1 was essentially based on previously published reports (*3-5*) with modifications to assess covalent binding. Recombinant Potentiation of Neddylation (PONY) domain of DCN1 and DCN2 proteins were generated using an E.coli expression system. These proteins were then biotinylated using the EZ sulfo-NHS-LC-biotin system (Thermofisher) and eventually labeled with terbium (Tb) by combining with streptavidin terbium (Tb) cryptate in a buffer with 200 mM NaCl, 25 mM Tris, 0.5 mM DTT, and 0.05% Tween20. The assay was run in the presence of a FAM-labeled competitive non-covalent DCN1 inhibitor, DI-591. The buffer solution containing the labeled DCN1 or DCN2 protein and the labeled DI-591 compound were added directly to the wells of the microtiter plates containing increasing concentrations of CLY-124 ranging from 0.051 nM to 3000 nM. The binding of the compound to the PONY domain was determined by the inhibition of the interaction with the competing probe, measured as a reduction in the TR-FRET signal using a plate reader at 1 and 24 h after treatment with compound (final DMSO concentration of 0.1%). The ratio was normalized to the high (DCN ½ + FAM-probe) and low (DCN1/2 without probe) controls for a readout of % activity.

The kinact and KI measurements were determined, under slightly modified conditions to allow for continuous reads over 10 h to assess time dependent changes in the IC50. Calculations for kinact and KI were based on the following equations (*6, 7*), where the KM of the probe is 34.56 nM. The concentration of the probe is represented by S in the equation. Plates were read every 5 min up to 1 hr, every 15 min up to 5 hr, and every hour up to 10 hours.

$$\mathrm{IC_{50}(t)} = \mathrm{K_I}\left(1 + \frac{\mathrm{S}}{\mathrm{K_M}}\right) \cdot \left(\frac{2 - 2e^{-\eta_{\mathrm{IC_{50}}} \cdot k_{\mathrm{inact}} \cdot t}}{\eta_{\mathrm{IC_{50}}} \cdot k_{\mathrm{inact}} \cdot t} - 1\right) \text{with}$$

$$\eta_{\mathrm{IC_{50}}} = \frac{\mathrm{IC_{50}(t)}}{\mathrm{K_I}\left(1 + \frac{\mathrm{S}}{\mathrm{K_M}}\right) + \mathrm{IC_{50}(t)}}.$$

**Pharmacokinetics of CLY-124 in NBSGW mice**

Whole blood was collected using K2-EDTA tubes and placed immediately on ice and was centrifuged within 30 mins at 2000 rcf at 4°C for 10 mins to isolate plasma. Plasma samples were then snap frozen using dry ice and stored at -80°C. A liquid chromatography coupled with tandem mass spectrometry (LC-MS/MS) method was established to quantify CLY-124 in mouse plasma samples. PK parameters were determined using non-compartmental analysis (PhoenixTM WinNonlin, version 8.5).

**In vivo Xenograft NBSGW mouse model setup and studies**

In vivo studies were performed essentially as outlined in fig S12A. Specifically, 300K human $CD34^+$ cells from G-CSF-mobilized human peripheral blood mononuclear cells (PBMCs) were injected into the tail veins of *NOD.Cg-Kit$^{W\text{-}41J}$ Tyr + Prkdc$^{scid}$ Il2rg$^{tm1Wjl}$/ThomJ* (NBSGW) mice (JAX Strain No. 026622) on day 0. Twelve weeks later, mice were tested for human cell engraftment by staining for expression of murine or human CD45 in PBMCs by flow cytometry. To ensure that similar levels of human progenitor cells were engrafted among the mice, mice with less than one percent or greater than ten percent human cells ($huCD45^+$) making up their PBMCs were removed from the study. Mice were then randomized and dosed using vehicle, HU or indicated doses of CLY-124. Two studies were performed. In the first study, groups of mice were treated with 25 mpk BID CLY-124, 100 mpk BID CLY-124, 50 mpk BID HU or the respective vehicles namely 5% Cremophor RH 40/20% hydroxypropyl-b-cyclodextrin for CLY-124 and sterile PBS for HU. Twice-a-day treatments for the mouse groups commenced during week 13 post engraftment and continued for 21 days, at which time the mice were euthanized, and their bone marrows individually collected for subsequent analysis. Body weights of mice receiving all the treatments were recorded daily. Bone marrow cells were filtered through a 40 mm cell strainer and preincubated with Human TruStain FcX™ (Biolegend #422302) and anti- mouse CD16/32 Ab (Biolegend # 101320) and then stained with fluorochrome-conjugated anti-human and anti-mouse CD45 to assess engraftment efficiencies.

For HU–CLY-124 combination studies, a staggered dosing regimen was implemented wherein HU was administered in the morning and CLY-124 was administered approximately 8 hours later in the evening. Staggered dosing was required due to the incompatibility of HU with the CLY-124 dosing formulation, which precluded co-administration.

*Measurement of HBG1 and HBB mRNA expression.* Bone marrow cell pellets (total 500K cells) were lysed in 25 µL of RLT (QIAGEN, 79216) with 1x β-mercaptoethanol (Gibco, 21985-023) and water followed by hybridizations. Hybridizations were done in a total volume of 15 µL (3 µL of RNA lysate added to master mix of 12 µL probe A/B (purchased from IDT), capture probe /reporter probes (NanoStringTM, XT TagSet-24, 121000602), proteinase K (Fisher Scientific, EO0491) and $H_2O$ suspended in hybridization buffer. HBG1 and HBB probe sets are listed in Supplementary Table 2. Samples were hybridized at 65 °C for 22 hr. Following hybridization, the tripartite complexes were purified, immobilized by nCounter Prep Station and imaged by Digital Analyzer (nCounter MAX/FLEX Analysis System), to generate digital counts of barcodes corresponding to each target in the multiplexed reaction. Labeled barcodes obtained from unamplified extracts were counted at 555 images or field of view (FOV). The barcode counts for each sample were recorded in Reporter Code Count (RCC) files that were imported into nSolver analysis software (provided with CodeSet by NS) for quality control evaluation.

*Synergy analysis for NBSGW HU and CLY-124 co-treatment study.* For assessment of pharmacologic synergy, a two-way analysis of variance (ANOVA) was performed with HU (HU) dose (vehicle, HU 75 mg/kg) and CLY-124 dose (vehicle, 5 mg/kg, 12.5 mg/kg) as fixed factors, including an interaction term (HU x CLY-124). HBG1 / (HBG1 + HBB) percentage was used as the dependent variable. Model residuals were visually inspected using residuals vs. fitted plots and Q-Q plots to confirm homoscedasticity and normality. Interaction term significance was used as the formal test for synergy. Analysis was performed using statsmodels (v0.14.4).

**Cytospin and histopathological characterization**

Cells from terminal erythroid cultures on day 14 were collected and cell suspensions made with 20,000 cells in 100µL of PBS + 0.5% BSA. Cells were then centrifuged to glass slides using a cytospin device (Thermo Scientific Shandon 4 Cytospin) at 900rpm for 5 min. Slides were air dried for 1hour and stained for 5 min in May-Grunwald stain (Sigma, MG-500) and rinsed in PBS for 1.5 min. Immediately after, slides were stained with Giemsa stain (1:20 dilution) (Sigma GS-500) for 20 minutes. Slides were rinse in deionized water and air dried. Cover slips were mounted using a drop of SecureMount (Fisherbrand Cat. No 022-208). Cell enumeration and scoring based on morphological features were performed by an ACP certified pathologist (Histowiz, NY).

**10x sequencing**

Human mPB $CD34^+$ cells were expanded and differentiated as described in the section “Cells and Erythroid Differentiation” and collected at day 0, day 3, day 5, and day 7 of differentiation (**fig S1A**). Cells were expanded for two days before the first treatment on day -2, which was two days prior to the start of erythroid differentiation. DMSO control, 33 nM MLN4924, 100 nM MLN4924, 100 nM DI-1548, or 3 µM DI-1548 were added to the culture following our standard treatment schedule described above. For CRISPR studies, electroporation was performed at day -2, and sequencing experiments included conditions containing electroporation only control or electroporation with RNPs targeting the enhancer of BCL11A, CUL3, UBE2M, and DCUN1D1 as described above.

*Sample preparation for single-cell RNA-sequencing*.
Harvested cells were stained using BioLegend TotalSeq-B anti-human hashtag antibodies B0251 to B0258 (BioLegend, 394631) on ice for 30 minutes. Each sample processed received a unique hashtag antibody, enabling pooling of multiple samples for downstream processing. After incubation, cells were washed three times to remove unbound antibody. Cells were then counted, and equivalent numbers of cells from all samples were pooled together into a new tube. Single-cell emulsions were generated using the Chromium Controller (10x Genomics, Pleasanton, CA, USA) with the Chromium Next GEM Chip G (10x Genomics, 1000120) and the Chromium Next GEM Single Cell 3' v3.1 Kit (10x Genomics, 1000121), targeting 12,000 cells per pool each day following the manufacturer's protocol. Briefly, cells and reverse transcription master mix were partitioned into oil droplets along with gel beads containing a poly(dT) primer for mRNA capture and a unique cell barcode sequence. Within droplets, cells were lysed enabling reverse transcription, producing barcoded, full-length cDNA from cellular mRNA. Hashtag-associated oligos were simultaneously captured by the gel bead incorporating the same cell barcode. Following reverse transcription, cDNA was amplified, and mRNA-derived cDNA was separated from hashtag oligo cDNA by SPRIselect cleanup and size selection (Beckman, B23318). Amplified cDNA products were evaluated on the Agilent BioAnalyzer 2100 using the High Sensitivity DNA Kit (Agilent, 5067-4626) for appropriate size and concentration, falling within 500 bp to 10 kb and greater than 40 ng/µL of cDNA. cDNA products were used as input for library construction incorporating Illumina adapters following manufacturer's instructions. Libraries were assessed for mass concentration using the Qubit 1X dsDNA HS Assay Kit (Invitrogen, Q33231) and the Qubit 4 Fluorometer (Invitrogen, Q33238). Library fragment size was evaluated using the Agilent BioAnalyzer 2100 using the High Sensitivity DNA Kit (Agilent, 5067-4626), with libraries passing QC between 400 and 2,000 bp. Libraries were then sequenced on the Illumina NovaSeq 6000 platform targeting a read depth of 50,000 read pairs per cell for gene expression libraries, and 10,000 reads per cell for hashtag libraries.

*Bioinformatics processing for single-cell RNA-sequencing*. Alignment, filtering, barcode error correction, and unique molecular identifier (UMI) counting were performed using the *count* function in the 10X Genomics software, Cell Ranger (v5.0.1). A custom GRCh38 (hg38) reference genome was used for alignment, and the introns were not included in the analysis. Hashtags were counted as separate features and appended to the count matrix. The merged count matrix was then subjected to a custom hashtag demultiplexing function that uses a multivariate gaussian mixture model to assign high confidence hashtag identities. Library-hashtag identities were then matched back to sample information.

**Single-cell RNA sequencing data analysis**
*Cell filtering and annotation*. Single cells and genes were filtered using functions in scanpy (v1.10.4) (*8*). All sequencing batches were integrated using scvi-tools (v1.3.0) to create a single reference embedding (*9*). Clusters were computed using the Leiden algorithm implemented in scanpy and individual cell types were labelled using cell type markers detailed in **fig. S1F**. Gene set activity scores were computed using the score_genes function in scanpy.

*Pseudotime analysis*. Pseudotime was computed using Palantir (v1.4.0) (*10*). In brief, this process includes calculation of a diffusion map, selection of a starting cell, construction of a Markov chain to determine the transition probability of individual cells and lastly computing the pseudotime as the average number of steps to reach each cell from the start cell, using the Markov process. For this analysis, an approximate starting cell was determined by searching for the cell with the highest CD34 expression in the HSPC cluster.

*Identifying differences in cell type composition.* To determine whether differences between control and perturbed conditions were likely to be biological rather than due to random sampling, we performed differential cell type composition analysis using py-scProportionTest (v0.1.2) (*11*). For each comparison between a control group (DMSO) and treatment group, we calculated the $\log_2$ fold difference in the proportion of each cell type and assessed significance using a permutation test (n=1000 permutations). Bootstrapped 95% confidence intervals (n=1000 iterations) were computed for observed differences, and multiple hypothesis testing was corrected using the Benjamini–Hochberg method.

*Cell cycle analysis.* To classify cell cycle states at single-cell resolution, we implemented a marker-based scoring strategy using curated gene sets for G0 (*'CDKN1B', 'CDKN2A', 'RB1', 'BTG2', 'GAS1', 'HES1'*), G1 (*'CCND1', 'CDK4', 'CDK6', 'MYC', 'RB1', 'CDKN1A'*), S (*'PCNA', 'MCM2', 'MCM4', 'MCM5', 'MCM6', 'MCM7', 'RPA1', 'RPA2', 'TYMS', 'BRCA1'*), and G2/M (*'CCNB1', 'CCNB2', 'CDK1', 'CDC20', 'PLK1', 'BUB1', 'AURKB', 'TOP2A'*) phases. For each cell, we computed the mean expression of phase-specific marker genes, extracted from either the primary expression matrix or a specified layer of the AnnData object. Unlike scanpy's score_genes_cell_cycle method, which scores S and G2/M phase markers relative to a randomly sampled background set and assigns each cell to S, G2/M, or otherwise G1 by exclusion, our method directly scores all four phases, G0, G1, S, and G2/M, and assigns the highest-scoring label per cell. We then applied a softmax transformation across phase scores to enable probabilistic interpretation and derived a confidence score based on the margin between the top two softmax-normalized values. This approach allows for more nuanced phase assignment, including identification of G0-arrested or G1-enriched cells, and captures uncertainty in classification.

*Signature scoring.* To characterize the transcriptional identity of the aberrant cell population, we computed per-cell gene-signature scores for four curated gene sets using the scanpy.tl.score_genes function, which calculates the mean expression of each gene set relative to a random background of comparable genes. The four signatures were: NRF2 pathway (*NQO1, TXNRD1, SQSTM1, FTH1, FTL, TXN, GSTO1*), cell cycle inhibition (*CDKN1A, CDKN1B, CDKN2A, CDKN2B*), mast-associated (*KIT, TPSAB1, TPSB2, TPSG1, GATA2, MITF, FCER1A*), and myeloid-associated (*CSF1, APOC1, CST3, S100A10, LAPTM5, CCRL2, TRPV2*). To avoid pseudoreplication inherent in single-cell data, signature scores were aggregated to pseudobulk estimates by computing the mean score per biological replicate within each cell type. Cell-type-level means were then calculated by averaging across all pseudobulk replicates per cell type. To enable comparison across signatures with different score magnitudes, values were Z-scored independently for each signature across the 11 cell types tested (HSPC, GMP, MEP, EryP, Ery, EBMP, EoP, aberrant, MkP, MonoP, MastP).

*Differential expression analysis.* We formed pseudobulked replicate samples, by summing the counts across all cells within a given sample. Only genes expressed in more than 5% of cells were considered for differential gene expression (DGE). To identify DGE, DMSO or EP Only reference groups were compared corresponding compound and genetic perturbations using Limma and defined at the thresholds $\log_2$(Fold-Change) >=1 and adjusted P-value < 0.05. For comparisons with cross sequencing library and/or cross-batch replicates, these fields were included as co-variates. To estimate the reproducibility of differential gene expression in response to our various CRISPR perturbations at day 0 and day 7 of erythroid differentiation, we performed a nonparametric bootstrap analysis on pseudobulked samples. In each of 100 bootstrap iterations, three biological replicates were sampled with replacement from each of the control (EP Only) and CRISPR KO groups and DGE analysis was performed.

**ATAC-seq sample preparation and sequencing**
Human mPB $CD34^{+}$ cells were expanded and differentiated as described in the section "Cells and Erythroid Differentiation". Cells were expanded for two days before the first treatment on day -2, which was two days prior to the start of erythroid differentiation. DMSO control, 100 nM MLN4924, or 100 nM DI-1548 were added to the culture following our standard treatment schedule described above. DMSO and treated cells were collected on day 5 of differentiation.

*Sample preparation for ATAC-sequencing*. ATAC-Seq libraries were prepared using the manufacturer's protocol (Active Motif, 53150) with minor modifications. Briefly, cells were collected from culture and counted, with 50,000 cells transferred to a new tube for ATAC-Seq processing. Cells were then washed once in PBS before being resuspended in cold lysis buffer and incubated on ice for 10 minutes. After lysis, nuclei were pelleted and resuspended in tagmentation mix with the transposase reaction carried out for 40 minutes at 37°C. The resulting tagmented DNA was purified, collected, and used as input for library construction incorporating Illumina sequencing adapters following manufacturer's instructions. After PCR cleanup, libraries were evaluated on the Agilent BioAnalyzer 2100 using the High Sensitivity DNA Kit (Agilent, 5067-4626) for a final library size ranging from 200 bp to 1,000 bp with a periodic peak each 150 bp. Libraries were assessed for mass concentration using the Qubit 1X dsDNA HS Assay Kit (Invitrogen, Q33231) and the Qubit 4 Fluorometer (Invitrogen, Q33238). Libraries were then sequenced on the Illumina NovaSeq 6000 platform targeting a read depth of 200M read pairs per library.

*Bioinformatics processing for ATAC-sequencing*. ATAC-seq libraries were processed using the nf-core/atacseq pipeline (v2.1.2, https://github.com/nf-core/atacseq/), which implements a standardized workflow for quality control, alignment, and peak identification. Adapter sequences were trimmed using Cutadapt (v3.4), and reads were aligned to the reference genome with BWA-MEM (v0.7.17). PCR duplicates were removed using Picard MarkDuplicates (v3.0.0), and reads were filtered using SAMtools (v1.17) to exclude unmapped and low-quality alignments. Signal tracks were generated as normalized bigwig files using BEDTools (v2.30.0) and deepTools (v3.5.1). Peaks were identified using MACS2 (v2.2.7.1) and genomic annotation of peaks was performed using HOMER (v4.11).

**CUT&RUN sample preparation, sequencing**
Human mPB $CD34^{+}$ cells were expanded and differentiated as described in the section "Cells and Erythroid Differentiation". Cells were expanded for two days before the first treatment on day -2, which was two days prior to the start of erythroid differentiation. DMSO control or 100 nM DI-1548 was added to the culture following our standard treatment schedule described above. Cells were collected at day 7 of differentiation. On the day of collection, cells were FACs sorted, to enrich the erythroblast population with FITC labeled anti-CD235 (BioLegend #306610) and APC labeled anti-CD71 (BioLegend #334108) antibodies, using a FACS sorter (Sony MA900).

*Sample preparation for CUT&RUN*. Following cell expansion and differentiation, cells were collected and counted. 10M cells per condition were washed once in PBS, then resuspended in 2mL nuclei extraction buffer (Epicypher, 21-1026). Lysis was carried out on ice for 10 minutes, then nuclei were resuspended in 1mL nuclei extraction buffer, counted, and frozen. All CUT&RUN sample processing was performed at EpiCypher (CUTANA™ platform, www.epicypher.com) using the manufacturer's kit and instructions. For each CUT&RUN reaction, 500K nuclei in Nuclei Extraction Buffer were dispensed into individual wells of a 96-well plate. Cells were then immobilized onto Concanavalin-A beads (Con-A) and incubated overnight at 4°C with 0.5 µg of each antibody (IgG, H3K4me3, NFYA). Following incubation, pAG-MNase was added to each well and activated by incubation at 4°C for 2 hours. Following which, CUT&RUN enriched DNA was purified using Serapure beads at a 1.7:1 (Bead:DNA) ratio. Recovered DNA was quantified using PicoGreen and reactions were normalized to 5 ng DNA before preparing sequencing libraries as per supplier's instructions. All CUT&RUN steps were performed on Tecan Freedom EVO robotics platforms. Sequencing was performed targeting 5-7M mappable 2x50bp paired-end reads per reaction on an Illumina NextSeq2000.

*Bioinformatics processing for CUT&RUN*. Sequencing libraries were processed through a custom pipeline developed by EpiCypher. Reads were aligned to the hg38 reference genome using Bowtie2 (v2.2.5)(*12*). Multi-mapping reads, PCR duplicates, and regions overlapping the ENCODE DAC exclusion list (*13*) were removed prior to downstream analysis. Normalized coverage tracks were generated with deepTools (v3.5.1) (*14*) with RPKM normalization, a bin size of 20 bp, and smoothing over 100 bp. Peaks were called using MACS2 (v2.2.7.1) (*15*) and annotated with HOMER (v4.11).

**Differential Peak calling and peak visualization**

To perform differential peak analysis between conditions from ATAC-sequencing and from CUT&RUN, consensus peaks were called across conditions using the merge function in BEDTools (v2.31.1) (*16*) and using replicate-merged peaks as input. A count matrix was then computed using the BEDTools function mulicov. Lastly, differential peaks were called using Limma. Replicate merged bigwig files, the consensus peak BED file, and gene BED files were visualized using pyGenomeTracks (v3.9) (*17*). The ch38 gene BED file was generated from ensemble (Homo_sapiens.GRCh38.105.gtf). To enhance visualization of differential chromatin accessibility genome-wide, we generated Manhattan plots from ATAC-seq peak-level statistics. Genomic coordinates were extracted by parsing peak identifiers in the format chr:start-end, using the midpoint as the representative peak position. To enable continuous plotting across chromosomes, cumulative genomic coordinates were calculated by ordering chromosomes (chr1-chr22, chrX, chrY) and sequentially offsetting the positions of each chromosome by the maximum position of the previous chromosome plus a 1 Mb spacer. Peaks were colored by chromosome and stratified statistically using a t-score threshold $t > 5$.

**Micro-Capture C sample preparation, sequencing, and analysis**

Human mPB $CD34^+$ cells were expanded and differentiated as described in the section "Cells and Erythroid Differentiation" and collected at day 7 of differentiation. Prior to collection, cells were FACs sorted to enrich erythroblast population with FITC labeled anti-CD235 (BioLegend #306610) and APC labeled anti-CD71 (BioLegend #334108) antibodies, using a FACS sorter (Sony MA900). Cells were expanded for two days prior to the first treatment on day -2, two days prior to erythroid differentiation phase, and treated with DMSO control or 100nM DI-1548 following our standard treatment schedule described above.

*Library preparation and sequencing*. All Micro-C library was prepared at Cantana Bio LLC using the Dovetail® Micro-C Kit (https://cantatabio.com/dovetail-genomics/), according to the manufacturer's protocol, on cells that were  cryopreserved in DMSO containing 10% FBS and sent to the service provider for processing. Each sample contained 1 million cells as starting input. Following thaw, the cells were fixed with disuccinimidyl glutarate (DSG) and formaldehyde. The cross-linked chromatin was then digested *in situ* with 0.5U micrococcal nuclease (MNase) per 1x10e6 cells. Following digestion, the cells were lysed with SDS to extract the chromatin fragments, and the chromatin fragments were bound to Chromatin Capture Beads. Next, the chromatin ends were repaired and ligated to a biotinylated bridge adapter followed by proximity ligation of adapter-containing ends. After proximity ligation, the crosslinks were reversed, the associated proteins were degraded, and the DNA was purified and then converted into a sequencing library using Illumina-compatible adaptors. Biotin-containing fragments were isolated using streptavidin beads prior to PCR amplification. The library was sequenced on an Illumina platform to generate at least 300 million 2 x 150 bp read pairs. Four libraries were constructed per sample targeting a total count of 1.2 billion reads per sample.

*Merging Pairs and Contact Matrix Generation.* Valid cis pairs (pairs occurring on the same chromosome) on chromosome 11 were filtered using *awk*. The resulting chr11-cis pairs were merged across replicates using *cat*. The experiments were normalized to the same number of valid pairs, based on the dataset with the lowest number of pairs (n = 59,081,584, 881 sample), with *shuf*. The normalized pairs were then sorted following the 4DN standard with the following command: *sort -k2,2 -k4,4 -k3,3n -k5,5n*. The resulting pairs were indexed using *pairix* (v0.3.7) and converted into a .cool file using *cooler cloud pairix* (v0.8.11) (*18*). A multi-resolution mcool file was generated at custom resolutions (500 bp, 1 kb, and 2 kb) using zoomify.
*Contact Matrix Resolution and Visual Assessments*. To determine the appropriate resolution for analysis, contact matrices were plotted over the region of interest (ROI) (chr11:5193824–5315567) at several resolutions (500 bp, 1 kb, and 2 kb). A read-support per bin analysis was also performed to identify the highest supported resolution. The pairs were converted into BED format using *awk*, and the bin structure from the .cool file was exported as BEDPE using the *cooler dump --join* command. The pairs were filtered using BEDTools (v2.28.0) with the *intersect -c* function to count the number of pairs per bin. Based on contact density and read support per bin, 1 kb was determined to be the highest resolution suitable for comparative analysis, as the read support per bin surpassed the requirement of >80% of bins containing a minimum of 1,000 reads (*19*).

*Relative Contact Probability (RCP)*. To validate whether detectable differences existed in contact profiles between the control (DMSO) and experimental condition (DI-1548) at the suggested resolution, a Relative Contact Probability (RCP) analysis was performed on chr11 at 1 kb resolution. The *RCP* function of GENOVA (v1.0.1) (*20*) was used to generate RCP curves and to calculate the log fold-change (logFC) of the RCP between DI-1548 and DMSO for each ATAC peak of within the ROI (N = 18 peaks). The RCP output was plotted using the visualize function in GENOVA. Detectable differences in the per-gene RCP curves further supported 1 kb as a suitable resolution.

*Virtual 4C and Differential Analysis*. Virtual 4C (v4c) was performed using GENOVA. Matrices were imported at 1 kb resolution and processed using the virtual_4C command for each matrix (DMSO and DI-1548) at every ATAC peak within the ROI. The results, per peak, were combined and contacts were dereplicated and exported in BEDPE format. Comparative analysis was performed with HicCompare (*21*). Briefly, the contacts were imported and joined with the *create.hic.table* function then normalized with *hic_loess*. Finally statistical differences were determined with *hic_compare* function with a threshold expression factor of 5, and an FDR p-value correction. Interactions with an FDR < 0.1 were identified as differential interactions and were exported as a BEDPE file. Significant interactions were dereplicated using mariner (v0.2.0) (*22*) with the *mergePair* function and a radius of 2x bin size (2,000 bp), as recommended by the tool.

**Statistical Analysis**

All statistical analyses were performed using GraphPad Prism unless otherwise stated in the Methods section.

**Supplementary Figures:**

**SUPPLEMENTARY FIGURE 1**

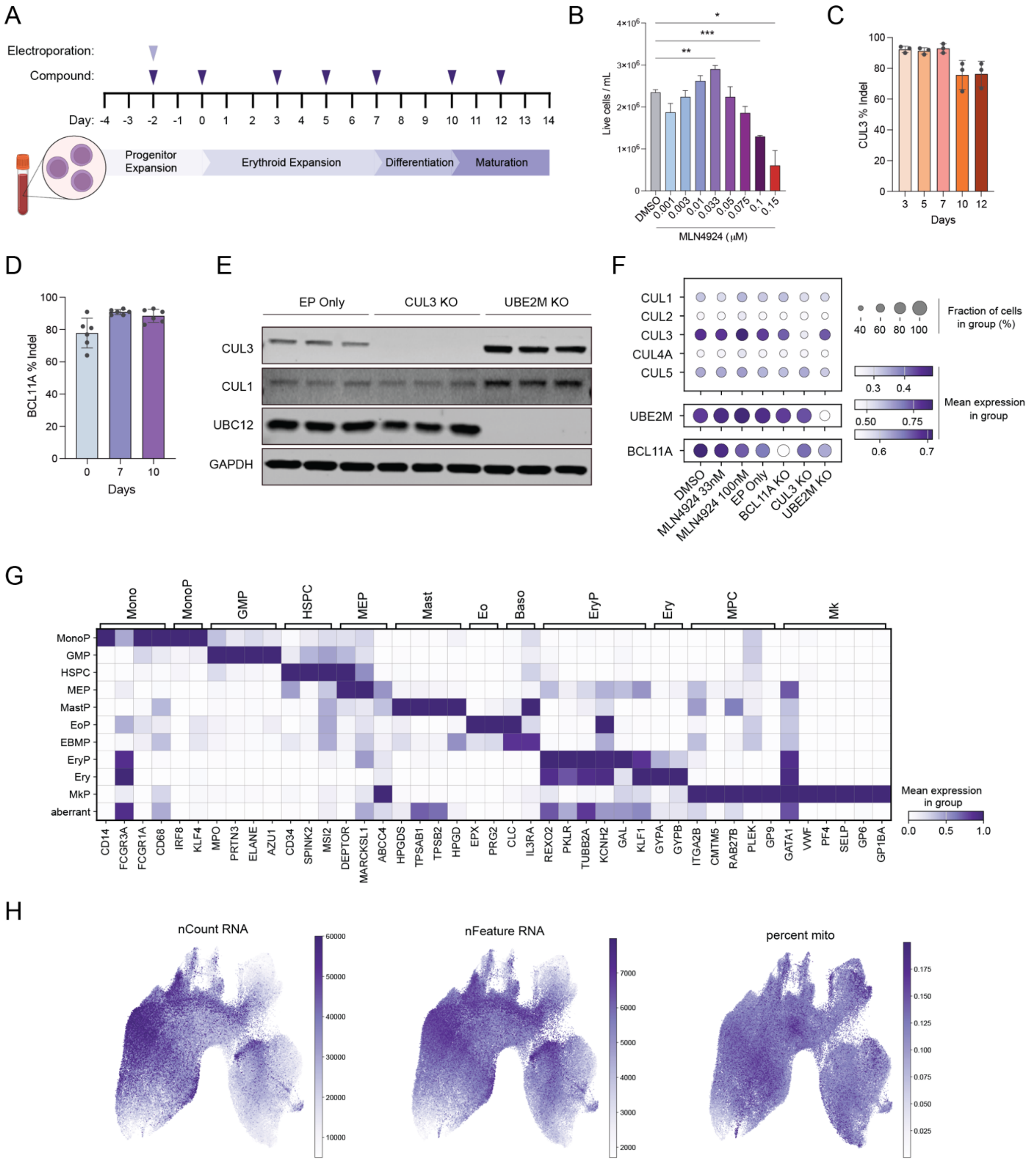


**Fig. S1. Quality control and experimental overview of CD34 cell culture and CRISPR perturbation in scRNA-seq**. **(A)** Schematic diagram of mobilized peripheral blood CD34 cell culture paradigm, including media changes, timepoints of compound additions (dark purple markers), and timepoints of electroporation of CRISPR guides (light purple marker). mPB CD34 image Created in BioRender. Miller, S. (2025) https://BioRender.com/nmmxtiw. (B) Significance determined by one-way ANOVA with Dunnett's multiple comparisons test (*p<0.05, **p<0.005, and

****p<0.0005). **(C)** and **(D)** Bar plot of CRISPR guide INDEL frequency over time. **(E)** Western blot validation of CRISPR knockouts. **(F)** Mean expression of target genes in scRNA-sequencing data at day 0 in HSPCs. (**G**) Matrixplot of scRNA-seq gene expression for markers of cell type lineages. **(H)** UMAP embeddings of standard QC metrics across all scRNA-seq timepoints.

### SUPPLEMENTARY FIGURE 2

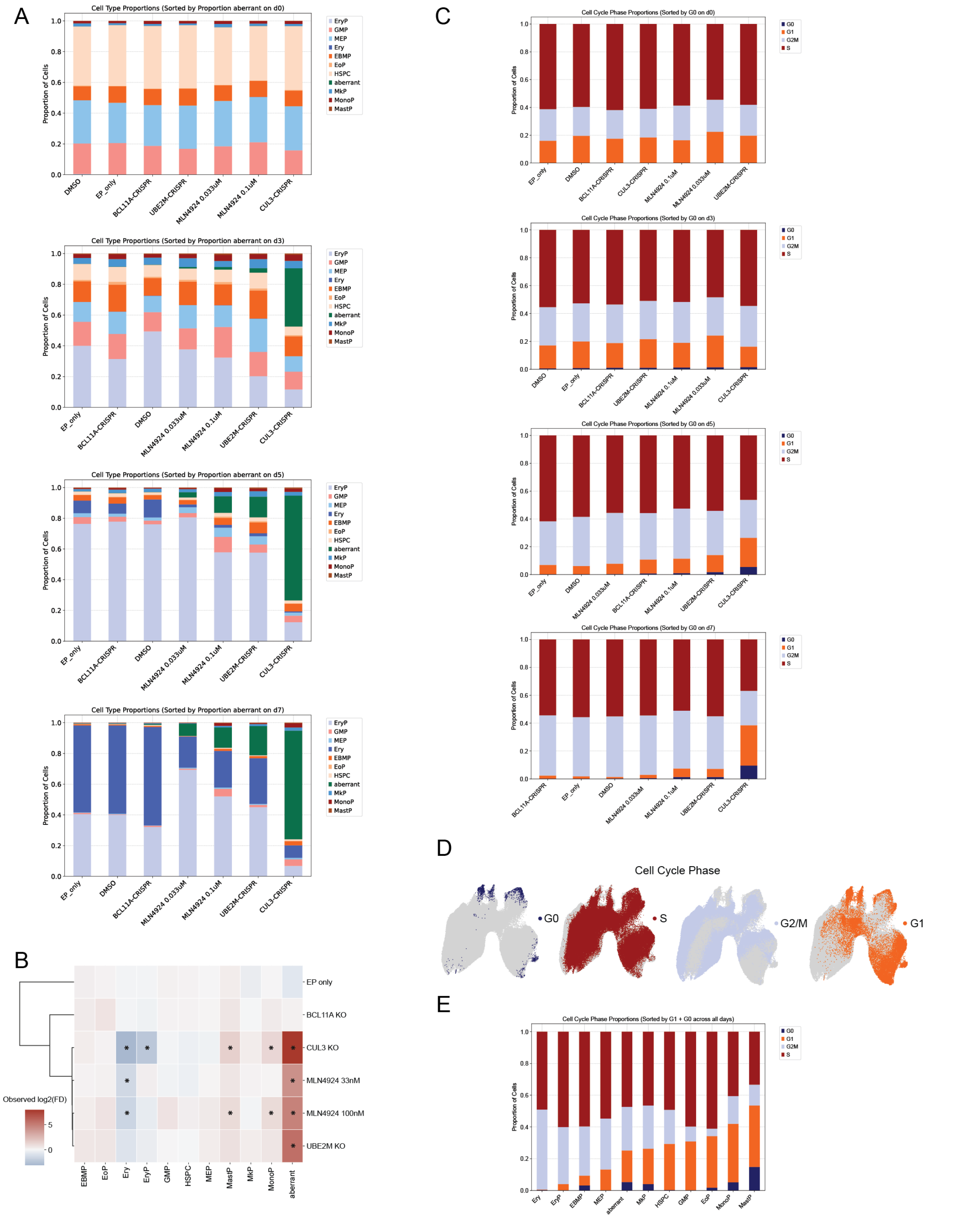


**Fig. S2. Neddylation pathway perturbations can alter cell type composition over time. (A)** Stacked bar plots showing the proportion of cells annotated for each cell type split by scRNA-seq timepoint. **(B)** Clustermap of scRNA-seq differential cell type enrichment results compared to DMSO computed across all timepoints combined and clustered by perturbation. (*) denotes $|log_2FD| \geq 1$ & adj $p < 0.05$. **(C)** Stacked bar plots showing the proportion of cells annotated for each inferred stage of the cell cycle split by scRNA-seq timepoint. **(D)** UMAP embeddings showing relative locations of cells annotated for each inferred cell cycle stage. **(E)** Stacked bar plots depicting the proportion of cells annotated for each inferred stage of the cell cycle across all scRNA-seq timepoints.

**SUPPLEMENTARY FIGURE 3**

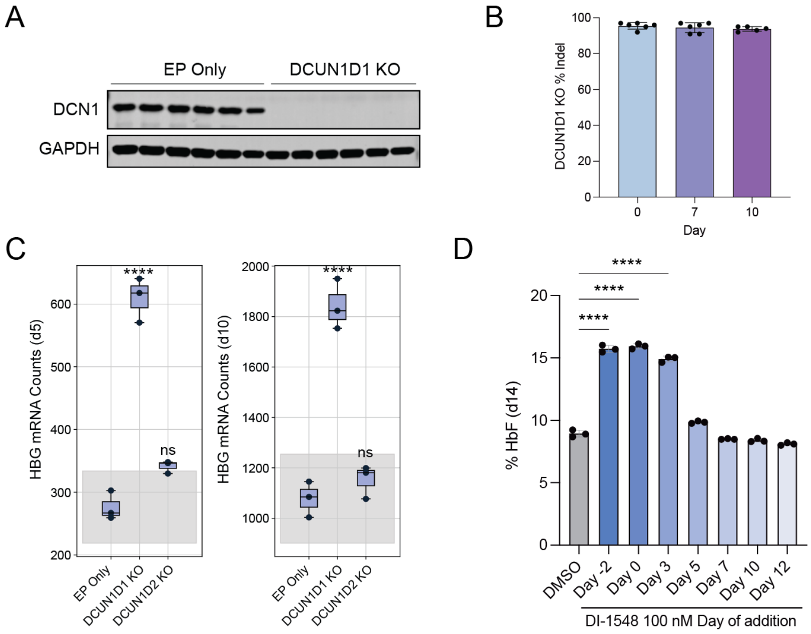


**Fig. S3. Targeting DCN1 but not DCN2 and timing of perturbation is critical for HbF induction. (A)** Western blot of DCN1 and GAPDH protein. **(B)** Bar plot of DCUN1D1 CRISPR guide INDEL percentage at days 0, 7, and 10. **(C)** Day 5 and day 10 normalized nCounter counts of HBG. Significance determined by one-way ANOVA with Dunnett's multiple comparisons test (****p<0.00005). Grey box indicates 95% confidence interval for the control condition, EP Only. **(D)** Day 14 readout of % HbF by HPLC after varying the starting timepoint of DI-1548 dosing. Significance determined by one-way ANOVA with Dunnett's multiple comparisons test (****p<0.0001).

**SUPPLEMENTARY FIGURE 4**

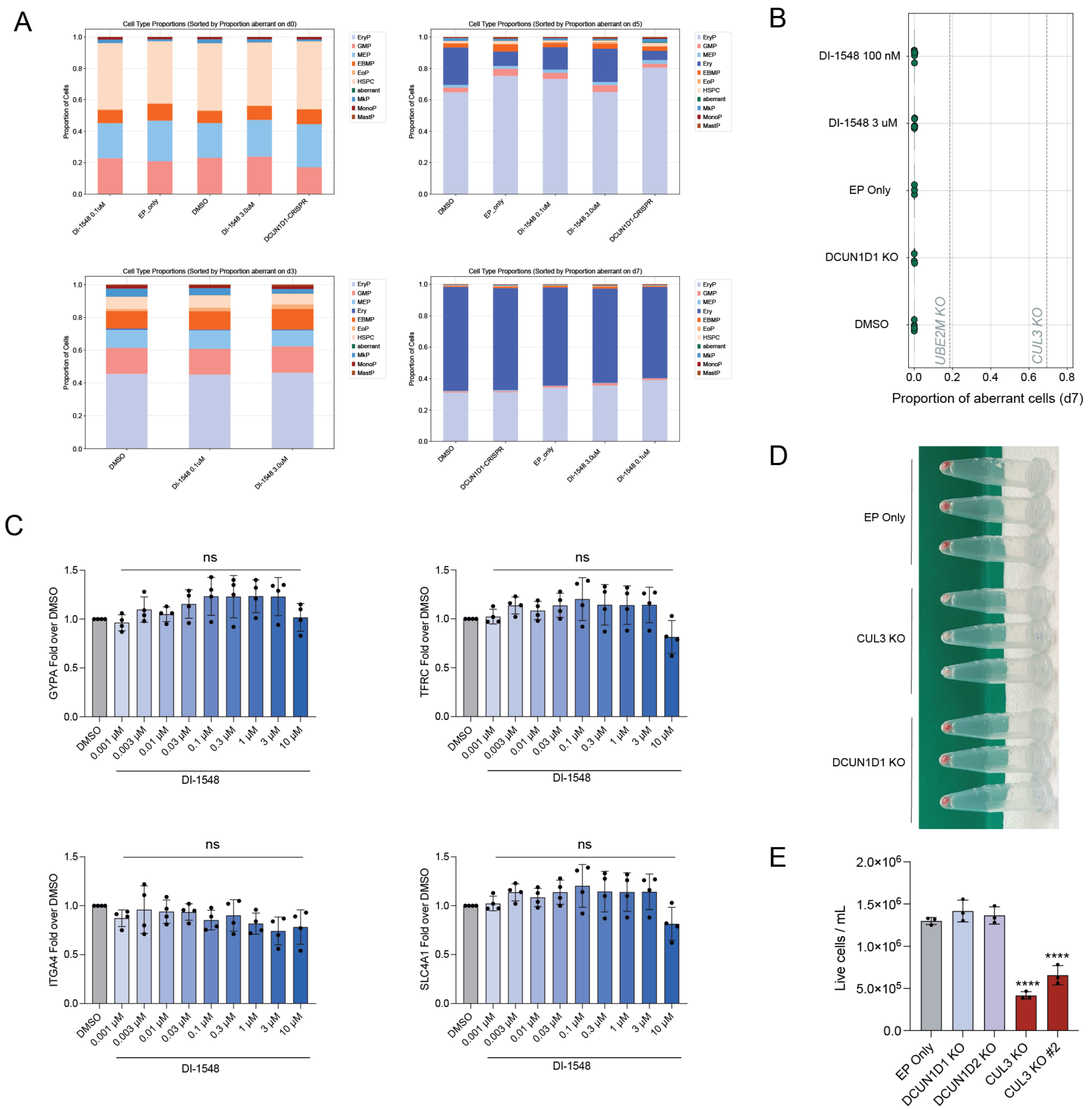


**Fig. S4. DCN1 inhibition does not impact erythropoiesis. (A)** Stacked bar plots depicting the proportion of cells annotated for each cell type split by scRNA-seq timepoint. **(B)** Bar plot showing the proportion of aberrant cells out of all cells on day 7 scRNA-seq. Statistical testing was not performed, as aberrant cells were negligibly rare across all conditions. **(C)** Relative fold change of various erythroid maturation marker gene expression across DI-1548 dose-response, measured by nCounter. Statistical significance determined by ANOVA with Dunnett's multiple comparisons test. **(D)** Images of cell pellets at differentiation day 14. **(E)** Live cell density on day 14 of erythroid differentiation.

### SUPPLEMENTARY FIGURE 5

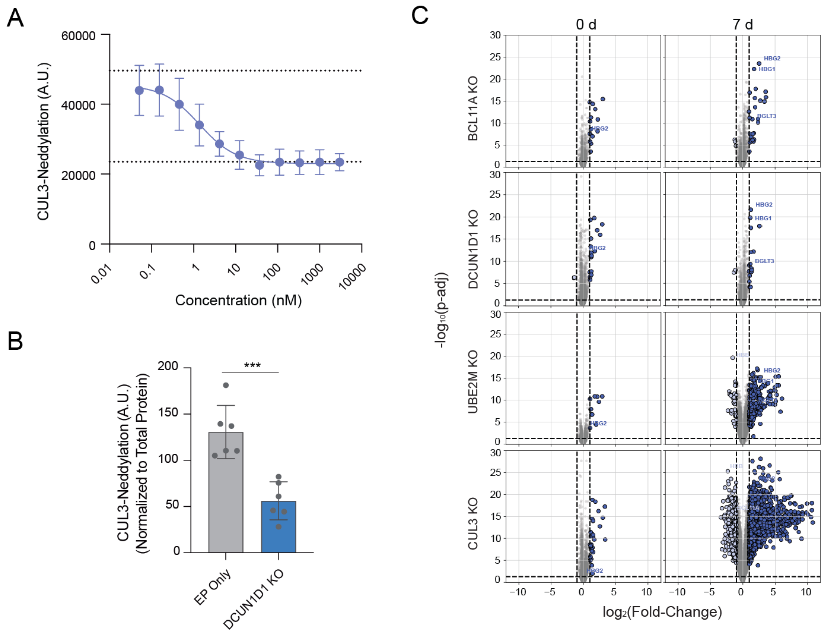


**Fig. S5. Inhibition of CUL3 neddylation and differential expression of neddylation pathway perturbations across timepoints. (A)** Inhibitory dose response curve of DI-1548 against CUL3 neddylation determined by AlphaLISA in TF-1 (N=7). **(B)** Inhibitory response against CUL3 neddylation determined by AlphaLISA. Significance was determined by unpaired t-test compared to EP Only (***p<0.0005). **(C)** Volcano plots of differential expression from replicates pseudobulked by timepoint with horizontal and vertical lines representing adj $p < 0.05$ and $|log_2FC| \geq 1$, respectively.

### SUPPLEMENTARY FIGURE 6

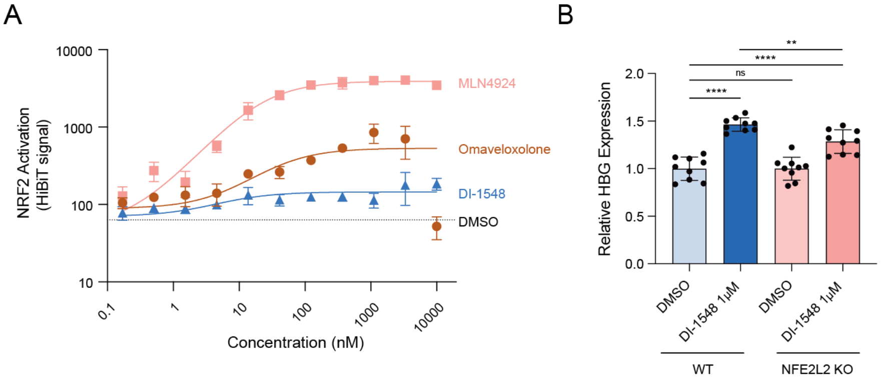


**Fig. S6. DCN1 inhibition modestly impacts BTB protein targeting of NRF2. (A)** Compound dose response curves of HiBiT-tagged NRF2 protein signal measured in HUDEP2 (N=4). **(B)** Relative fold change in HBG counts measured by nCounter after compound stimulation in wild-type (WT) or NRF2 KO HUDEP2 cells.

**SUPPLEMENTARY FIGURE 7**

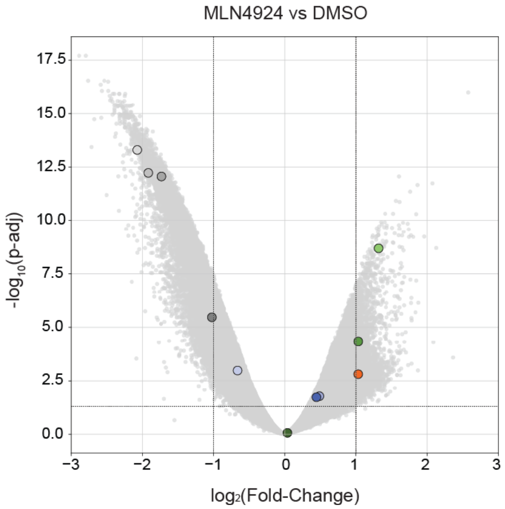


**Fig. S7. Impact on chromatin accessibility by neddylation pathway inhibitors.** Volcano plot of differential peaks (N=5). Dot color corresponds to peak color in Fig. 3B, with all other peaks colored grey.

**SUPPLEMENTARY FIGURE 8**

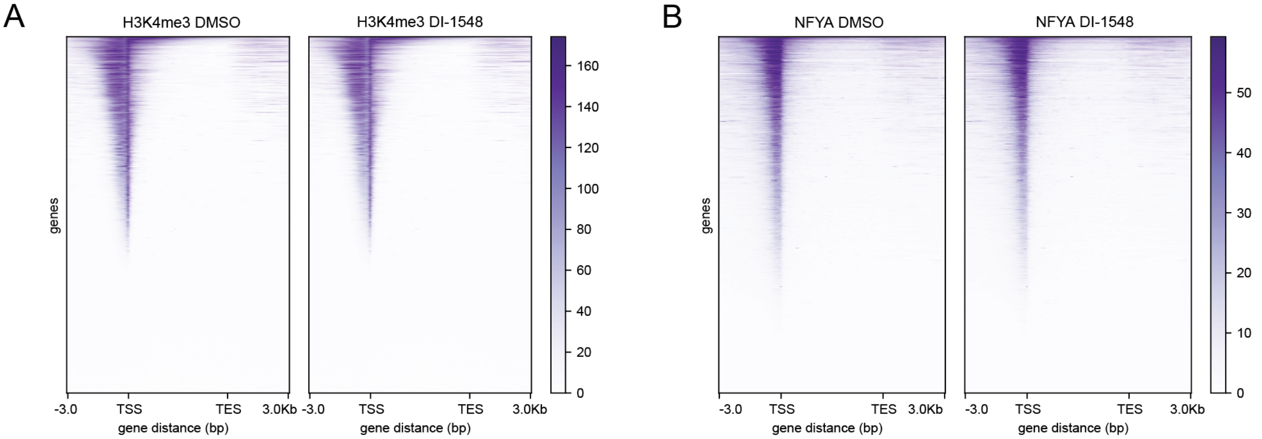


**Fig S8. H3K4me3 and NFYA show TSS enrichment pattern.** Enrichment heatmap for **(A)** H3K4me3, and **(B)** NFYA, across genes genome wide (n=3). TSS = transcription start site, TES = transcription end site.

### SUPPLEMENTARY FIGURE 9

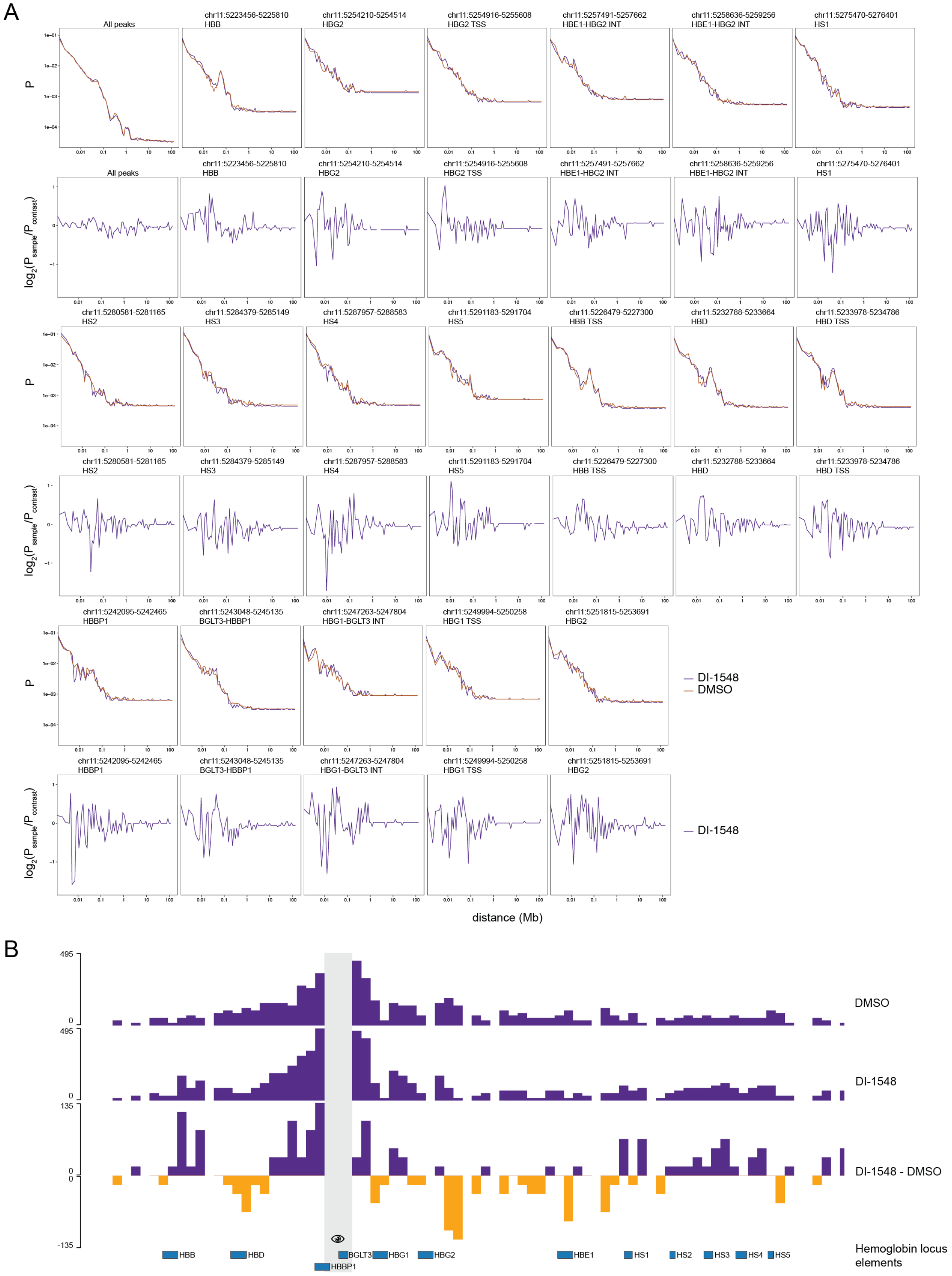

**Fig. S9. DCN1 inhibition alters the contact profile at the hemoglobin locus. (A)** Probability of MCC signal from anchor points moving away from selected genomic loci (n=2). **(B)** BED graphs of MCC signal or signal subtracted between DI-1548 and DMSO across the globin locus (n=2).

### SUPPLEMENTARY FIGURE 10

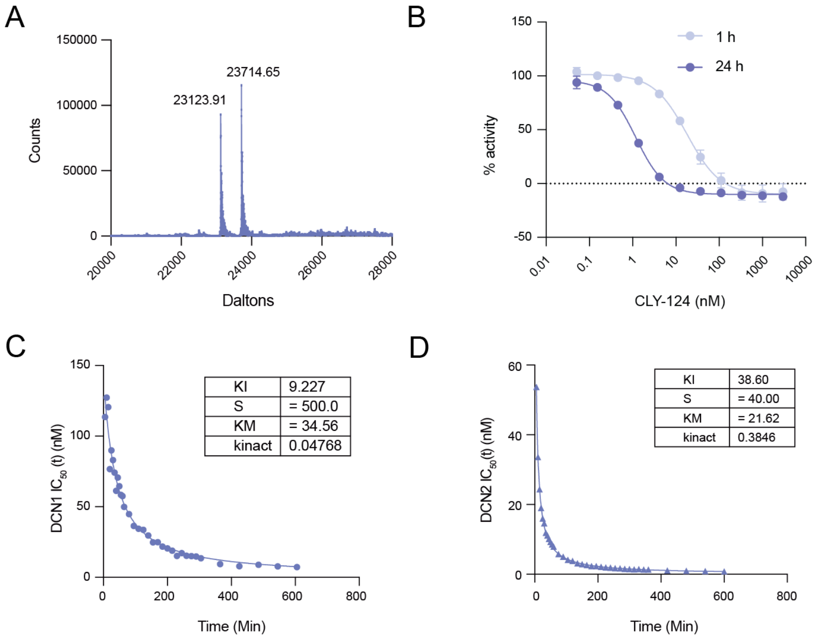


**Fig. S10. CLY-124 binds to DCN1 and DCN2. (A)** Representative mass spectrum of DCN1 (400 nM) incubated with CLY-124 (370 nM) for 3 hours, depicting peaks from both apo-protein (23123.91 kDa) and CLY-124:DCN1 adduct (23714.65). **(B)** Dose response analysis of CLY-124 competitive binding to DCN1 protein at 0.31 nM using TR-FRET assay in the presence of a FAM-labeled probe, DI-591 at 100 nM and unlabeled probe at 800 nM, after 1 and 24 hours of incubation (n=4 for each time point). **(C)** Estimation of $k_{inact}$ and KI of CLY-124 binding to DCN1 protein (final conc 0.31 nM) from IC50 values derived over time. Measurements shown here were obtained from total probe concentration of 500 nM. KI and KM are in nM units and $k_{inact}$ is in $min^{-1}$. **(D)** Estimation of $k_{inact}$ and KI of CLY-124 binding to DCN2 protein (final conc 0.31 nM) from IC50 values derived over time. Measurements shown here were obtained from total probe concentration of 40 nM. KI and KM are in nM units and $k_{inact}$ is in $min^{-1}$.

**SUPPLEMENTARY FIGURE 11**

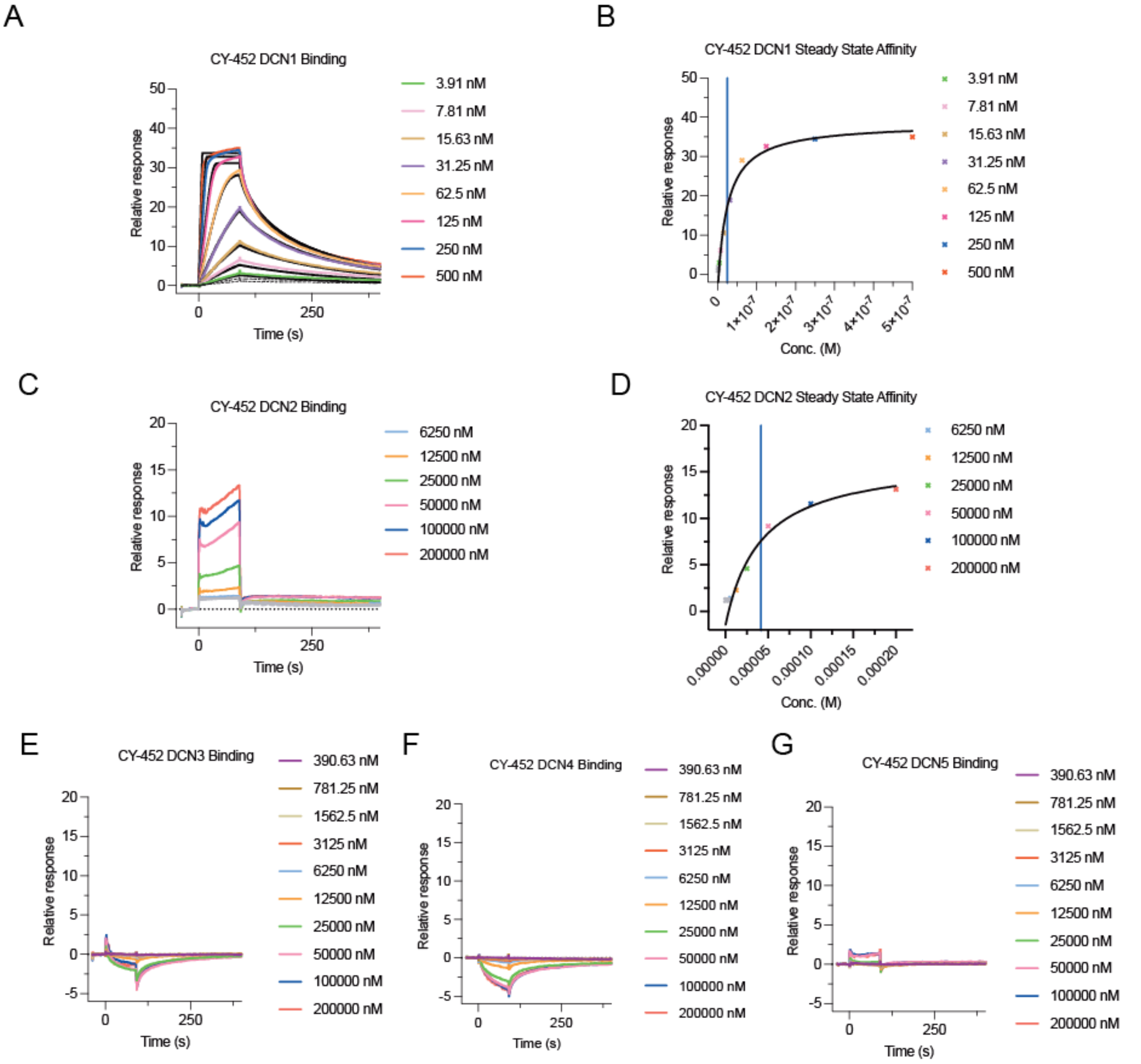


**Fig. S11. SPR analysis of CLY-124 binding to DCN isoforms 1, 2, 3, 4, and 5. (A)** Sensorgram depicting binding of CLY-124 to DCN1. **(B)** Representative SPR versus concentration plot from steady state binding of indicated concentrations of CLY-124 to DCN1; Blue line indicates KD determined from the binding. **(C)** Sensorgram depicting binding of CLY-124 to DCN2. **(D)** Representative SPR versus concentration plot from steady state binding of indicated concentrations of CLY-124 to DCN2; Blue line indicates KD determined from the binding. **(E)** Sensorgram depicting lack of binding of indicated concentrations of CLY-124 to DCN3, **(F)** DCN4, and **(G)** DCN5.

### SUPPLEMENTARY FIGURE 12

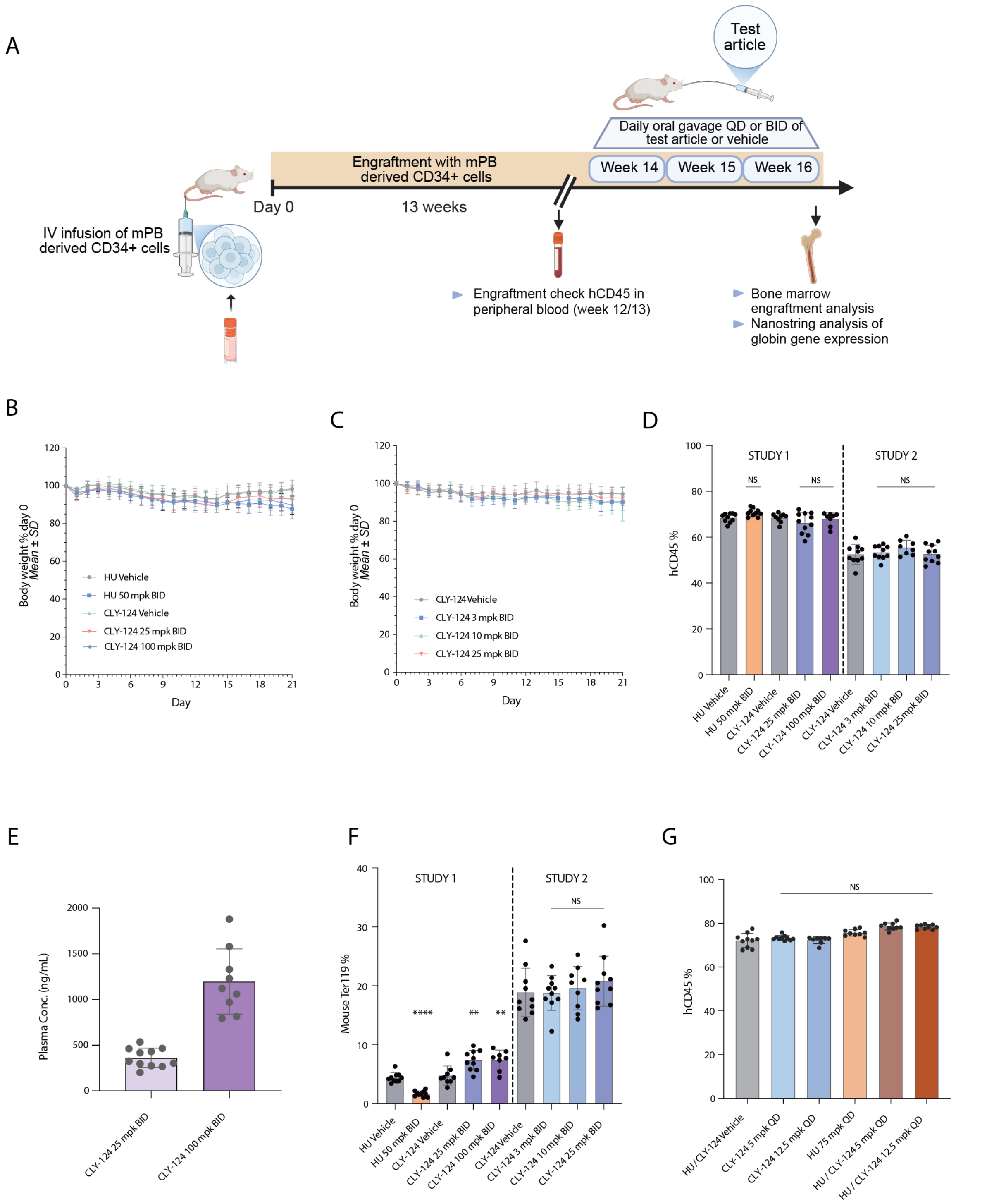


**Fig. S12. CLY-124 in-vivo dosing shows no impact on body weight, mouse erythropoiesis, and engraftment efficiency. (A)** Schematic of NBSGW mouse studies outlining human CD34$^+$ cell engraftment, testing for engraftment efficiency, compound administration, and sample harvesting for analysis. Created in BioRender. Krishnamoorthy, S. (2025) https://BioRender.com/wzab91q. **(B)** and **(C)** Body weights of mice dosed with indicated interventions over time for 21 days plotted as percentage of day 0 (pre-dose), data plotted as mean ± SD. **(D)** Percentage of engrafted cells in the NBSGW mice determined by flow cytometry as % $hCD45^+$ cells. **(E)** CLY-124 exposure in mouse plasma measured after 21 days of dosing with 25 mpk and 100 mpk BID. Data presented as mean ± SD of plasma concentration in ng/mL. **(F)** Flow cytometry analysis of murine $Ter119^+$ cells in bone marrow of mice after administration of indicated compounds for 21 days. Data plotted as mean ± SD of % $Ter119^+$ bone marrow cells from each mouse. **(G)** Percentage of engrafted cells as in (D). For (D, F-G) statistical significance was determined using unpaired t-test or one-way ANOVA with Dunnett's multiple comparisons test, as appropriate (**p<0.005, ****p<0.00005).

**Supplementary Tables:**

**SUPPLEMENTARY TABLE 1**

| | |
|---|---|
| **In vitro Screening** | |
| Assay | |
| DCN1 TR-FRET (nM / Geometric Mean) | |
| 1h | 80.84 */ 2.66 nM |
| 24h | 4.98 */ 1.70 nM |
| $k_{inact}$/Ki | |
| DCN1 | 0.004647 nM-1*min-1 |
| DCN2 | 0.0025 nM-1*min-1 |
| KD from SPR Analysis | |
| DCN1 | 13.9 nM |
| DCN2 | 41.5 mM |
| DCN3, 4, 5 | No binding |
| TF-1 cell target occupancy (IC50) | 10.15 nM |
| CUL3-NEDD8 AlphaLISA (IC50) | 15 nM |
| **In vitro ADME** | |
| MW I ALogD I pKa | 590 \| 5.1 \| 10.7 |
| Blood Stability - T1/2 (min) (human I rat I mouse I dog I monkey) | 245 \| 41 \| 30 \| >427 \| 208 |
| % remaining at 120min | 57 \| 8.1 \| 3.0 \| 79 \| 55 |
| Hepatic Stability - T1/2 (min) (human I rat I mouse I dog I monkey) | 94 \| 65 \| 42 \| 176 \|68 |
| Clint (ml/min/10e6 cells) (human I rat I mouse I dog I monkey) | 14.8 \| 21.4 \| 32.9 \| 7.92 \| 20.5 |
| PPB fu (%) (human I rat I mouse I dog ) | 0.75 \| 0.95 \| 0.58 \| 0.47 |
| Blood to plasma partition (human I rat I mouse I dog ) | 0.64 \| 0.49 \| 0.76 \| 0.57 |
| Caco-2 Permeability and Efflux (x 10-6cm/s, recovery) | A->B 1.71 \| 60% |
| | B->A 5.10 \| 78% |
| | ER 3.0 |
| Thermodynamic Solubility in Bio-relevant media (mg/ml) (FaSSIF **\|** FeSSIF) | 25 \| 77 |
| Thermodynamic Solubility in PBS pH 7.4 (mg/ml) | 3.79 |

**Table S1. In vitro screening and ADME characteristics of CLY-124**

**SUPPLEMENTARY TABLE 2**

| Gene | Accession No. | Probe A | Probe B |
|---|---|---|---|
| HBB | NM_000518.4 | AATTGGACAGCAAGAAAGCGAGCTTAGTGATACTTGTGGGCCAGGGCATTCATCCTCTTCTTTTCTTGGTGTTGAGAAGATGCTC | CGAAAGCCATGACCTCCGATCACTCATCCCCCAGTTTAGTAGTTGGACTTAGGGAACAAAGGAACCTTTAATAGA |
| HBG1 | NM_000559.2 | GTTCTCAGGATCCACATGCAGCTTGTCACAGTGCAGCTGTTGAGATTATTGAGCTTCATCATGACCAGAAG | CGAAAGCCATGACCTCCGATCACTCCCGAAATGGATTGCCAAAACGGTCACCAGCACATTTCCCAGGAGCTTGAA |
| GYPA | NM_002099.4 | TTCCTCTCTCTTCATCTTACGCTTTTGCTTGCCTTGCTGCTTTGTTATCACGAACCTAACTCCTCGCTACATTCCTATTGTTTTC | CGAAAGCCATGACCTCCGATCACTCCGCCTTATAGAGGGAATGGGCTTCTGATGAAAGGATGTGTTTGGCTTCAT |
| TFRC | NM_003234.1 | GTCTGGACAGGTATATTAGGCAATCCTGATGACCGAGATGGTGGAAACTGCCAATTTGGTTTTACTCCCCTCGATTATGCGGAGT | CGAAAGCCATGACCTCCGATCACTCAGTCTCCTTCCATATTCCCAAACAGCTTTTCTGCAGCAGCTCTGGAGATT |
| ITGA4 | NM_000885.4 | TCGCCCCGGGATTGATCACTGAAGCGTTGGCGAGCCAGTTGCTTTCGGGTTATATCTATCATTTACTTGACACCCT | CGAAAGCCATGACCTCCGATCACTCCTGTTCGCACGTCTGGCCGGGATTCTTTCCGATCCTGCATCTGTAAA |
| SLC4A1 | NM_000342.3 | AGGAGGCCGCCGAAGGTGATGGCGGGTGACAGTGCAGCAACAACAGCCACTTTTTTTCCAAATTTTGCAAGAGCC | CGAAAGCCATGACCTCCGATCACTCCAGCTCCGACACTCCCATCTGGTTCCGGGTCTTTTCTCCC |
| POLR2A | NM_000937.2 | ACTGGCCCAACAGGAAGACAGTAAGCGAAGGAGTCTTTGGCTTCTTGGAACGAACCTAACTCCTCGCTACATTCCTATTGTTTTC | CGAAAGCCATGACCTCCGATCACTCCAGACGGCACAGAATATCCTTGGCTCTCTCAGCATCTCGAGCGG |
| SDHA | NM_004168.1 | TAAACCCTGCCTCAGAAAGGCCAAATGCAGCTCGCAAGCCTGCCCAATTTGGTTTTACTCCCCTCGATTATGCGGAGT | CGAAAGCCATGACCTCCGATCACTCTGCAACAGTGTGTGACCTGGTAGGAAACAGCTTGGTAACACATGCTGTAT |
| GUSB | NM_000181.3 | CTTTTTATTCCCCAGCACTCTCGTCGGTGACTGTTCAGTCATGAAATCGGCTTTCGGGTTATATCTATCATTTACTTGACACCCT | CGAAAGCCATGACCTCCGATCACTCCGCAAAAGGAACGCTGCACTTTTTGGTTGTCTCTGCCGAGTGAAGATCCC |
| RPL19 | NM_000981.3 | AATCCTCATTCTCCTCATCCATGTGACCTTCTCTGGCATTCGGGCATTGGCAACAGCCACTTTTTTTCCAAATTTTGCAAGAGCC | CGAAAGCCATGACCTCCGATCACTCTGGCGATCGATCTTCTTAGATTCACGGTATCTTCTGAGCAGCCGGCGCAA |
| ABCF1 | NM_001090.2 | CCAGCTTGATGTCAGATGCATTTTCTAACATGGCTTGGCGGGAGGACATCCCTGGAGTTTATGTATTGCCAACGAGTTTGTCTTT | CGAAAGCCATGACCTCCGATCACTCGTCTGCATTGACGAACAGCTCCTTGCCATGAGCGGAGATGCTGAACTTCT |

| | | | |
|---|---|---|---|
| G6PD | NM_000402.2 | CTCAGTGCCAAAGGGCTCCTTGAAGGTGAGGATAACGCAGGCGATGTTGTCAGATAAGGTTGTTATTGTGGAGGATGTTACTACA | CGAAAGCCATGACCTCCGATCACTCTGCATCACGTCCCGGATGATCCCAAATTCATCGAAATAGCCCCCGCGACC |

**Table S2. Nanostring targets and probe sequences**

**SUPPLEMENTARY TABLE 3**

| Antibody | Fluorochrome | Vendor | Catalog number | clone |
|---|---|---|---|---|
| CD71 | PE | BD Biosciences | 566722 | OKT9 |
| CD235a (GlyA) | PE-CY7 | Biolegend | 306620 | HIR2 |
| CD49b (integrin a4) | Brilliant Violet 421 | BD Bioscience | 565277 | 9F10 |
| CD233 (Band 3) | FITC | IBGRL | 9439 | BRIC6 |
| CD45 | PerCP | BD Biosciences | 347464 | 2D1 |
| HbF | FITC | Life Technologies | MHFM01-4 | HBF-1 |
| NFYA | Unlabeled | Bethyl Laboratories | #A302-105A | Polyclonal |
| H3K4Me3 | Unlabeled | Epicypher | #13-0060 | 2909-3D7 |
| IgG | Unlabeled | Epicypher | #13-0042 | Polyclonal |

**Table S3: List of antibodies used for flow cytometry**

**SUPPLEMENTARY TABLE 4**

| Gene Name | gRNA Sequences |
|---|---|
| BCL11A (+58 Enhancer) | CTAACAGTTGCTTTTATCAC |
| CUL3 | ATCCAGCGTAAGAATAACAG |
| CUL3 #2 | GGTGTATTAGGGATCATCTA |
| UBE2M | GCGCAGCTGCGGATCCAGAA |
| DCUN1D1 | ACAGCAGTTCTGTGATGACC |
| DCUN1D2 | TCAGTGTATTGGTCATAGCG |
| NFE2L2 | GGUUUCUGACUGGAUGUGCU |

**Table S4: Sequences of gRNAs**
